# PIFiA: Self-supervised Approach for Protein Functional Annotation from Single-Cell Imaging Data

**DOI:** 10.1101/2023.02.24.529975

**Authors:** Anastasia Razdaibiedina, Alexander Brechalov, Helena Friesen, Mojca Mattiazzi Usaj, Myra Paz David Masinas, Harsha Garadi Suresh, Kyle Wang, Charles Boone, Jimmy Ba, Brenda Andrews

**Author notes:** Department of Chemistry and Biology, Toronto Metropolitan University, Toronto, ON, Canada.

## Abstract

Fluorescence microscopy data describe protein localization patterns at single-cell resolution and have the potential to reveal whole-proteome functional information with remarkable precision. Yet, extracting biologically meaningful representations from cell micrographs remains a major challenge. Existing approaches often fail to learn robust and noise-invariant features or rely on supervised labels for accurate annotations. We developed PIFiA, (**P**rotein **I**mage-based **F**unct**i**onal **A**nnotation), a self-supervised approach for protein functional annotation from single-cell imaging data. We imaged the global yeast ORF-GFP collection and applied PIFiA to generate protein feature profiles from single-cell images of fluorescently tagged proteins. We show that PIFiA outperforms existing approaches for molecular representation learning and describe a range of downstream analysis tasks to explore the information content of the feature profiles. Specifically, we cluster extracted features into a hierarchy of functional organization, study cell population heterogeneity, and develop techniques to distinguish multi-localizing proteins and identify functional modules. Finally, we confirm new PIFiA predictions using a colocalization assay, suggesting previously unappreciated biological roles for several proteins. Paired with a fully interactive website (https://thecellvision.org/pifia/), PIFiA is a resource for the quantitative analysis of protein organization within the cell.

## Introduction

Recent progress in high-throughput microscopy and computational image analysis has catalyzed large-scale efforts to quantitatively describe single-cell biology^1-6^. Advances in quantitative analysis of large-scale image datasets have been driven by the development of algorithms for protein localization prediction, which have been used for automated drug screening, and extracting morphological profiles from cell images^7, 8^. Computational methods enable efficient analysis of millions of single-cell images by extracting morphological information in an unbiased quantitative form. However, generating meaningful numerical features from single-cell images remains a significant challenge. Cells in micrographs typically exhibit a variety of shapes and positions, while noise levels and pixel intensities can also vary between images, making it difficult to develop algorithms that extract functionally rich patterns while ignoring irrelevant information^8^. For instance, early machine learning approaches relied on hand-engineered feature sets extracted from images, such as cell texture and shape, which were often difficult to select and not transferable to other datasets or tasks^2^. Ideally, a computational workflow would map single cells and proteins to robust numerical representations, enabling analysis of the spatial organization of the cell in an objective way.

More recently, single-cell images have been analyzed using deep learning methods, which overcome the limitations associated with hand-engineered feature sets by learning the optimal feature representations directly from pixel level data^2,9^. Of particular relevance, we previously developed a deep convolutional neural network, DeepLoc^8^, for analysis of images of GFP-fusion proteins for each budding yeast open reading frame (ORF). The yeast ORF-GFP collection^10^ consists of ∼4,100 unique strains, each of which expresses a GFP signal above background in standard growth conditions, enabling systematic analysis of ∼70% of the yeast proteome in living cells^2, 8, 10^. DeepLoc was trained to accurately classify images from diverse datasets into 22 subcellular compartments, including images generated in different genetic backgrounds and by different laboratories^8, 11^. Similar approaches have been developed to analyze human cell datasets, such as the Human Protein Atlas, a collection of immunofluorescence images which covers ∼65% of the human proteome^12^. While supervised approaches produce high-quality features, their success largely depends on the number of hand-labeled samples in the dataset^13, 14^. For example, the Human Atlas Project leveraged crowd-sourcing to accelerate label collection on a large scale, involving thousands of video games players for image annotation^12^. However, manual label assignment is not practical for imaging datasets containing millions of single-cell micrographs. In addition, human-labeled standards may reflect the biases of an individual annotator and can preclude identification of subtle or incompletely penetrant phenotypes^14, 15^.

An emerging alternative to supervised methods for biological image analysis involves self-supervised approaches, which do not require manually assigned categories during training^16-18^. Instead, self-supervised learning models define a training objective, or pretext task, using structural information from the data itself^18-20^. In the context of self-supervised training, features learned with the pretext task should encapsulate information from the images that is useful for downstream applications, such as the discovery of common localization patterns by clustering analysis^15^. Recently, self-supervised methods based on auto-encoders have been used for representation learning on cellular data^21-23^.

Autoencoder-based models are trained by compressing an image into the latent space (encoding), and subsequent image reconstruction (decoding)^24^. The encoding of the image in the latent space is then used as its representation. For instance, Paired Cell Inpainting^15^, a self-supervised approach developed for analysis of yeast fluorescent micrographs, encodes several imaging channels to predict the appearance of a fluorescently-tagged protein in a target cell. Another autoencoder-based method developed for human cell data, *cytoself*^*22*^, trains a vector-quantized variational autoencoder^24, 25^ (VQ-VAE) to reconstruct fluorescent signals of tagged proteins. Self-supervised learning with autoencoder-based approaches has also been applied for the analysis of human microglia data^21^, and extraction of feature profiles predictive of cell metastatic potential^23^. A disadvantage of autoencoders is that they often learn features that are not relevant to protein morphology and localization, such as cell position, imaging artifacts and noise^15^. Also, pixel-level reconstruction is computationally expensive and often unnecessary for representation learning. These issues reflect the autoencoder’s training objective, which targets identical reconstruction of the input. In this study, we asked whether other characteristics of microscopy data could be leveraged as self-supervised objectives to learn high-quality image representations.

Another challenge related to learning image-based features lies in their downstream analysis and interpretation. Current approaches typically extract representations with various machine learning methods and perform downstream analysis using clustering and tSNE/UMAP projections^8, 15, 22^. However, there are no clear rules for more nuanced biological analysis, including analysis of extracted features for different levels of cellular organization, or high-confidence identification of protein functional modules. In general, data-backed guidelines on hyperparameter selection, which enable biologically meaningful clustering and consider the scale of cellular organization, are needed. Also, current molecular representation learning approaches generally lack methodologies that can characterize protein function by quantifying cell-to-cell variability in individual protein behavior. In summary, a gap remains in the image analysis field, requiring approaches that could (1) learn biologically meaningful features without human annotations, (2) produce universal features useful for studying subcellular organization at different scales, and (3) provide techniques for a wide range of downstream feature analyses.

Here, we present PIFiA (**P**rotein **I**mage-based **F**unct**i**onal **A**nnotation), a self-supervised approach for protein functional annotation derived from single-cell imaging data. PIFiA is coupled with a range of feature exploratory techniques for biological discovery. The representation learning component of PIFiA is performed by a convolutional neural network (CNN), which was trained with the objective of predicting protein identity directly from its fluorescently-labeled input image. This objective does not depend on pre-existing annotations or human labels and, unlike autoencoder-based models, PIFiA is robust to learning non-relevant information in the image, such as cell position, multiple cells in a crop, input noise or imaging defects. In addition to the CNN component, the PIFiA workflow includes a set of downstream analysis steps for quantitative exploration of feature profiles extracted from single-cell imaging data. We applied PIFiA to ∼3,000,000 live-cell confocal images of the budding yeast open reading frame (ORF)-GFP fusion collection^10^. We compare PIFiA to existing approaches for protein representation learning and show that PIFiA outperforms previous methods on four different standards of protein function. We explore PIFiA feature profiles for use in a variety of downstream tasks, which are designed for the discovery of functional groups across different scales of cellular organization. Solely using distinct localization patterns of each protein, PIFiA can make remarkably precise functional predictions, identifying highly specific subcellular localization and distinct functional modules to reveal new biological insights.

## Results

### PIFiA architecture, feature profiles and proteome-scale image dataset

PIFiA is a self-supervised deep learning approach designed to derive functional information about proteins from microscopy data without using any pre-existing annotations (**Fig. 1**). The PIFiA workflow consists of a feature extraction step performed by a deep neural network (**Fig. 1a, b**), as well as subsequent analysis steps on the extracted feature profiles (**Fig. 1c, d, e**). The downstream analysis enables prediction of protein localization and the identification of functional modules or subsets of proteins with related cellular roles, such as protein complexes and their associated regulators. The feature profiles can be used for multiple downstream tasks, including construction of a hierarchical map of subcellular organization (**Fig. 1c**), predicting protein function (**Fig. 1d**), identifying localization heterogeneity at a cell population level (**Fig. 1e**), and finding functional modules.

**Figure 1.**
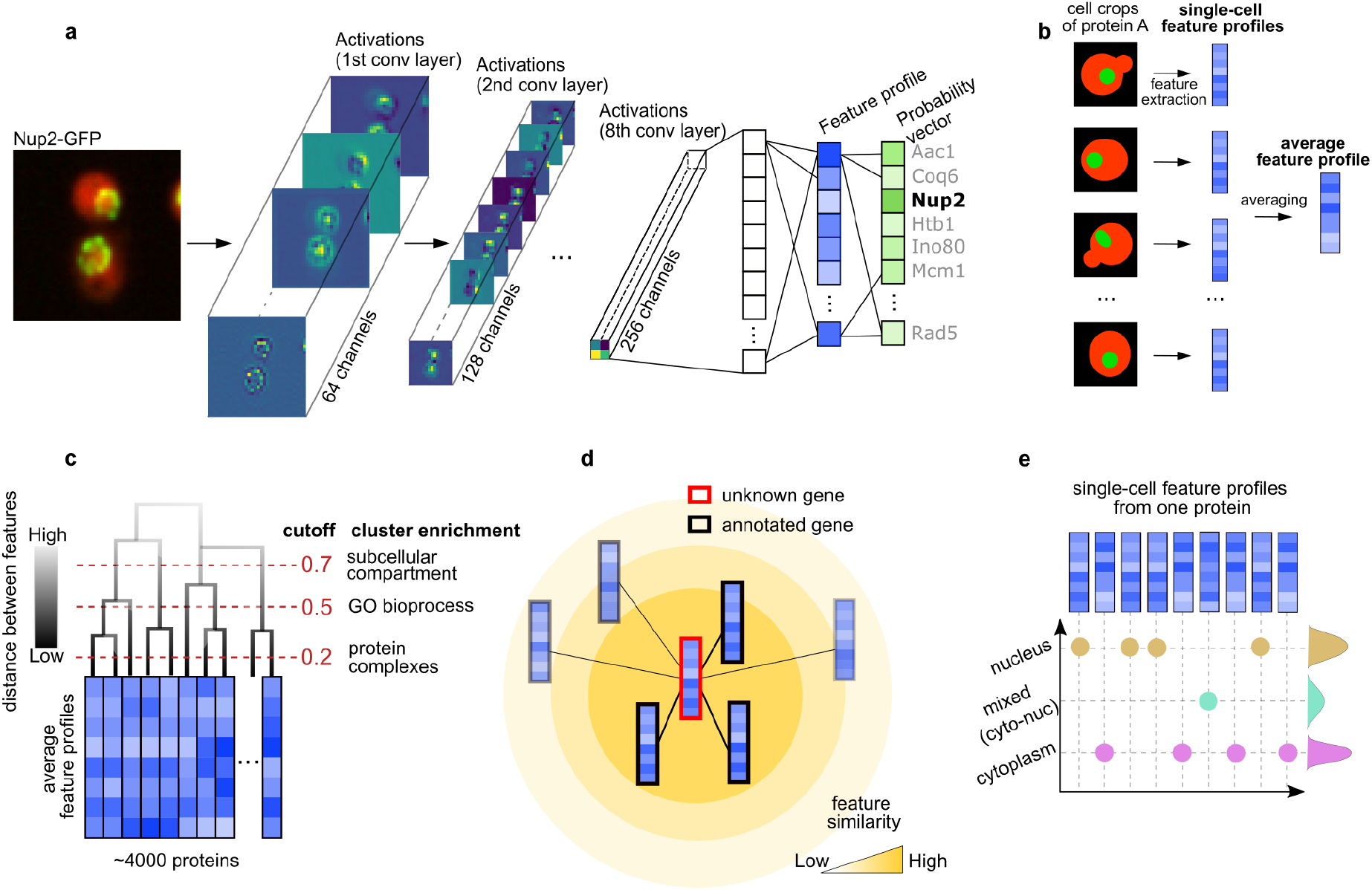
Overview of the PIFiA workflow. **a**, The backbone of PIFiA is a deep learning model, which contains convolutional blocks followed by fully-connected layers. Shown are examples of activations from passing a micrograph of fluorescently labeled Nup2 protein (Nup2-GFP) through the PIFiA network, with corresponding patterns recognized by the convolutional filters. Feature profiles are extracted from the second fully-connected layer, for use in downstream applications (c, d, e). **b**, Illustration of two types of feature profiles produced by PIFiA - single-cell feature profiles (extracted from a single crop) and averaged feature profiles (obtained by averaging all single-cell feature profiles of that protein). **c**, Schematic representation of the global hierarchy of protein feature profile similarities to reveal different levels of functional information. **d**, Illustration of protein function prediction using self-supervised PIFiA feature profiles. **e**, An illustrative example of using PIFiA single-cell feature profiles to investigate the localization heterogeneity of a protein.

The deep learning backbone of PIFiA is a CNN consisting of eight convolutional blocks and three fully-connected (FC) layers, which was trained to predict a protein identifier associated with an input image (i.e. one out of 4,049 classes (**Fig. 1a**)). The CNN produces a feature profile (or a representation profile) from the input image, which is unique to a particular image. Feature profiles are 64-dimensional real valued vectors extracted from the second FC layer, which is followed by a classification layer (**Fig. S1a**). These feature profiles encapsulate condensed information about each protein’s identity, based solely on its localization pattern. Over the course of training, the model first learns straightforward characteristics, such as patterns of different cellular compartments, then it subsequently learns more subtle morphological features that may distinguish individual proteins (**Fig. S1b**). To achieve the best accuracy and simplicity trade-off, we searched for the optimal architecture, network depth/width and related hyperparameters based on the validation set (**Fig. S1c, d**) (see Methods). We found that more complex architectures, such as DenseNets^26^, did not improve performance but increased the training time, hence we chose a simpler architecture that could achieve comparable performance. Similarly, we searched for optimal feature profile dimensionality and found that accuracy of protein identity prediction stabilized around a 64-dimensional feature profile (**Fig. S1d**).

To train PIFiA, we produced a comprehensive dataset of 3,058,961 live-cell images of individual strains expressing both a unique fusion gene from the yeast ORF-GFP collection^10^ and spatial markers of cell cycle position^10, 14, 27^, which provide cellular context for computational analysis of protein localization. In particular, we used automated yeast genetics^28^ to engineer a new version of the ORF-GFP collection, in which the resultant strains also carried fluorescent markers of the nucleus (td-Tomato-NLS) and cytoplasm (E2-Crimson). In total, images of 4,049 unique strains were obtained using an automated confocal microscope. Cell images were derived from two biological replicates, each of which had four fields of view for each ORF-GFP strain. The images acquired for the GFP channel were cropped into 64×64 pixels crops (median of 778 crops per tagged protein, see Methods), and each crop contained at least one cell at its center. The crops for each GFP-tagged protein were then split into training, validation and test subsets (8:1:1 ratios).

After CNN training was completed, we extracted feature profiles of the individual single-cell crops from the test set to produce single-cell feature profiles (scFPs) (**Fig. 1b**). We then averaged the scFPs for each protein to create its average feature profile (aFP) (**Fig. 1b**). An aFP and scFP for an individual protein have the same dimensions, but they describe different levels of information: scFPs encapsulate the localization pattern of one cell, while aFPs describe the general spatial distribution of a protein. Below, we first use PIFiA aFPs to broadly explore protein localization and function. We then use scFPs to explore cell-to-cell heterogeneity, localization changes, and complex protein localization patterns (**Fig. 1c-e**).

### Comparison of PIFiA performance to other self-supervised and supervised approaches

We compared aFPs produced by PIFiA to the representations from three computational methods, each of which has been used previously to analyze images of the yeast ORF-GFP collection^10^: CellProfiler^7^, a feature-extraction tool; DeepLoc^8^, a supervised deep-learning model; and Paired Cell Inpainting^15^, a self-supervised autoencoder-based approach. We consider a model to have good performance if protein pairs with higher correlation between their aFPs are more likely to be functionally related. We evaluated feature profiles (aFPs) using three metrics: F-score and average precision (AP), both measures of feature relevance, and adjusted mutual information (AMI), an information theoretic metric to assess clustering quality (see Methods). PIFiA features showed superior performance on most evaluation criteria using four functional benchmarks: Gene Ontology^29^ (GO) Cellular Components (CC), GO Slim Bioprocesses (GO BP slim), Kyoto Encyclopedia of Genes and Genomes^30^ (KEGG) pathways and European Bioinformatics Institute (EBI) Protein Complexes^31, 32^ (**Fig. 2**).

**Figure 2.**
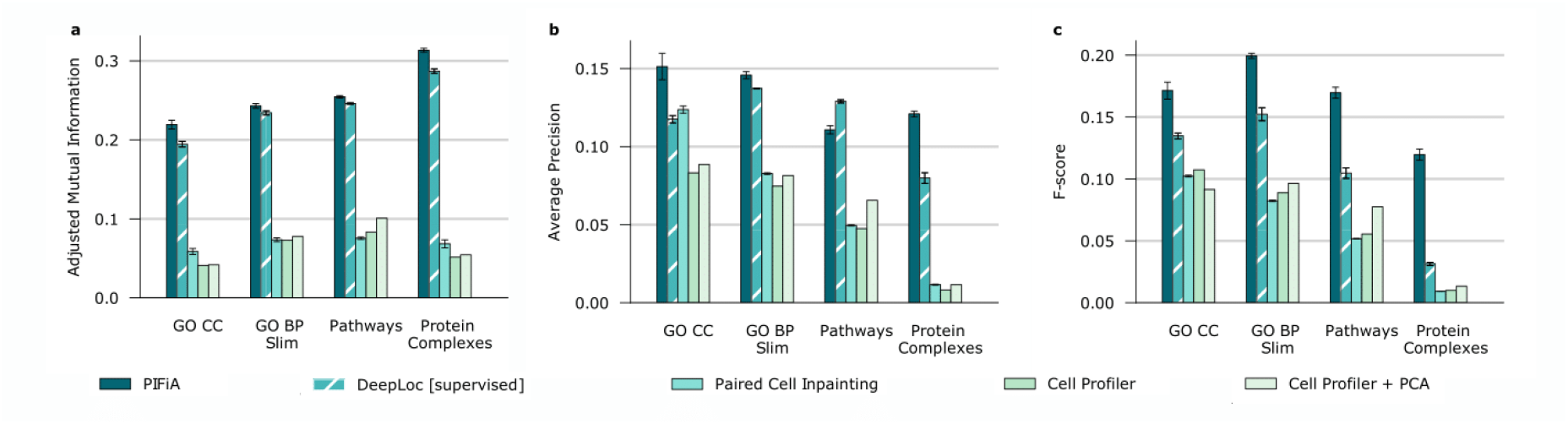
Comparison of PIFiA performance to the existing supervised and self-supervised methods for protein representation learning. Bar graphs show the performance of PIFiA and four other methods (X axis) at detecting pairs of functionally related genes/proteins. Performance was assessed using: **a**, the adjusted mutual information; **b**, average precision; **c**, F-score on four biological standards [X axis: Gene Ontology (GO) Cellular Component (CC), GO Slim Bioprocess (GO BP Slim), Kyoto Encyclopedia of Genes and Genomes Pathways (KEGG pathways) and European Bioinformatics Institute protein complexes (Protein Complexes). Error bars indicate the standard deviation of the scores across three independent runs (for deep learning models).

PIFiA reached better performance than the supervised method DeepLoc in predicting protein subcellular localization (DeepLoc’s target task), as indicated by higher values of F-score, AP, and AMI scores, on the GO CC standard. (**Fig. 2**). PIFiA also outperformed DeepLoc based on other functional standards, with the biggest performance gain in protein complex discovery. This result confirms the utility of the PIFiA training objective which targets identification of individual tagged proteins, the most detailed level of functional information present in the image. Although the objective does not directly focus on localization prediction, over the course of training the CNN implicitly learns a variety of localization patterns needed to successfully differentiate individual proteins. Thus, PIFiA self-supervised feature profiles can be used for exploratory analysis of protein localization instead of representations from a supervised method such as DeepLoc, bypassing the need for manual annotation while improving performance.

PIFiA also demonstrated better performance than Paired Cell Inpainting, another self-supervised method, achieving 1.2, 1.7, 2.2 and 10.4-fold improvements in terms of mean average precision using cellular components, bioprocesses, pathways and protein complex standards, respectively. Compared to all other approaches examined, PIFiA representations resulted in substantial improvement in clustering quality measured by AMI scores, with an average 5-fold AMI improvement over Paired Cell Inpainting (**Fig. 2a**). The significant improvement on the protein complex standard is again explained by PIFiA’s novel self-supervised objective, which forces the network to detect the most comprehensive morphological patterns while ignoring individual image artifacts, which contrasts with autoencoder-based objectives that learn features by naive image reconstruction.

### Evaluation of the functional information associated with PIFiA average feature profiles

To further assess the biological information associated with the aFPs of each protein, we used hierarchical clustering of aFPs as an unsupervised approach to discover feature profile similarities^33^. We performed agglomerative hierarchical clustering of the whole-proteome aFPs (4049 proteins in total) using a correlation metric and average linkage. We surveyed the resulting dendrogram at different thresholds to explore whether aFPs are suitable for studying the spatial architecture of the cell at different scales of its organization (**Fig. 3a, S2a-f**). The hierarchical clustering results are shown in **Fig. 3a**, with 4,049 proteins on the X-axis clustered according to the similarity of their feature profiles (each column is a 64-dimensional aFP). To determine optimal cutoff thresholds, we tracked AMI scores^34^ at different correlation thresholds for three functional standards: GO Cellular Component, GO Slim Bioprocess and Protein Complexes (**Fig. S2b**) (see Methods).

**Figure 3.**
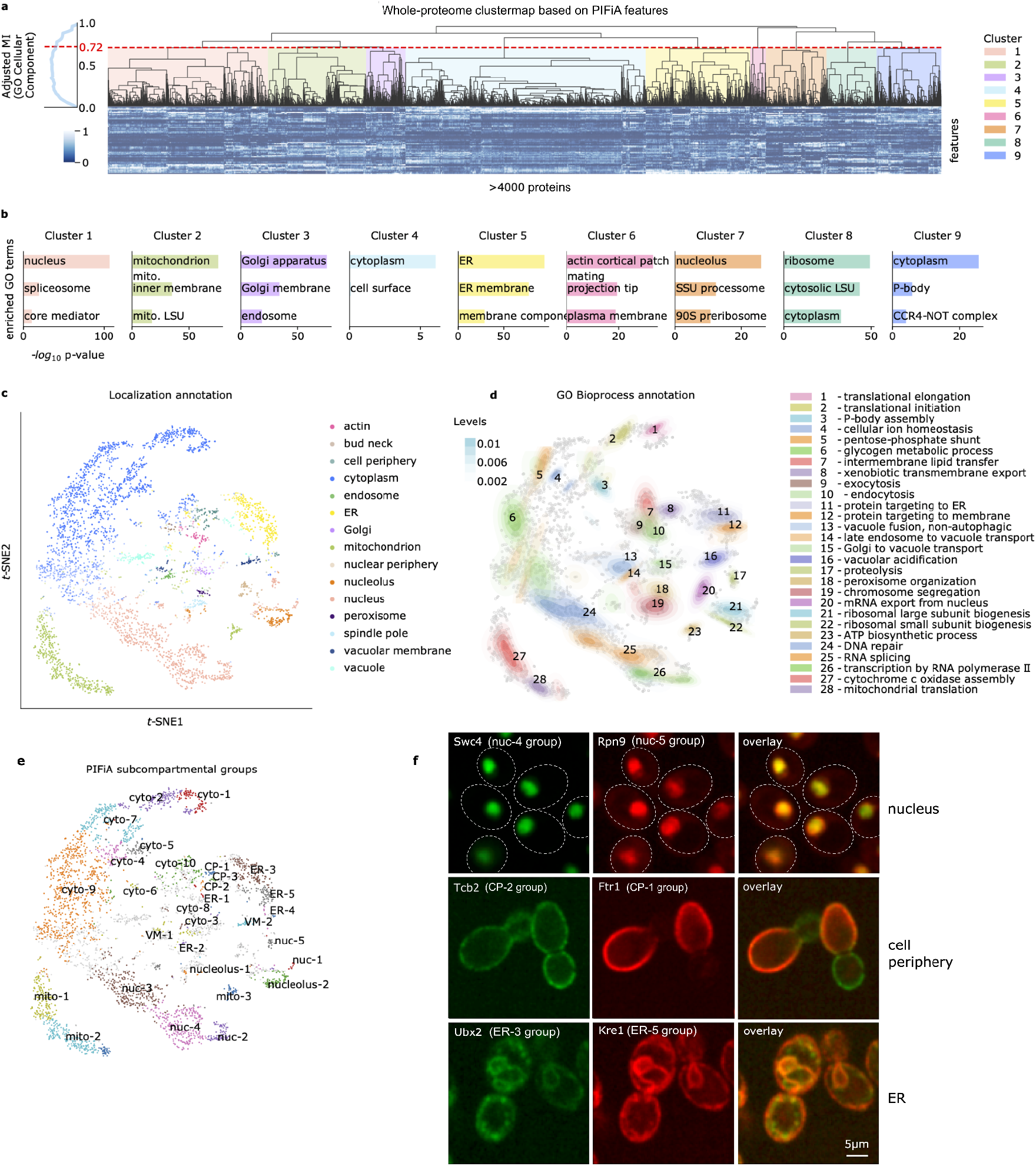
Clustering of PIFiA average feature profiles and analysis of the associated biological information. **a**, Subcellular organization revealed by clustering of PIFiA’s average feature profiles. The plot to the left of the Y axis shows the adjusted mutual information curve between clustering labels and GO Cellular Component labels at different distance thresholds. The distance threshold (d = 0.72) indicated on the clustergram produces clusters associated with cell compartments (color codes on the right). **b**, The top three Gene Ontology Cellular Component scores for each cluster defined in a are shown. **c**, Whole-proteome tSNE projection of PIFiA average feature profiles. Each point on the plot represents a protein colored according to a localization category predicted by logistic regression (see Results for training details). **d**, Annotation of the whole-proteome tSNE projection with GO bioprocess categories shown as Gaussian kernel density estimates. Bioprocesses were selected according to the lowest variance from different cellular components. The color intensity of the kernel density estimate contour plots corresponds to the cumulative probability mass below the drawn contour. **e**, Annotation of the whole proteome tSNE projection with sub-compartmental protein groups predicted by clustering of PIFiA feature profiles (cyto - cytoplasmic cluster, nuc - nuclear cluster, mito - mitochondrial cluster etc.; full descriptions of clusters are in **Table S2**). **f**, Representative micrographs of cells expressing mNeonGreen- (left; green images) or mScarlet- (middle; red images) tagged proteins annotated to different sub-compartmental groups within three cellular compartments: *nucleus* (top), *cell periphery* (middle) or *endoplasmic reticulum* (ER, bottom) groups. Overlays of the mNeonGreen and mScarlet images are shown on the right. The tagged proteins are indicated on the micrographs (scale bar shown bottom right).

First, we determined an optimal threshold (0.72) corresponding to the most general level of cellular organization - GO Cellular Component annotations (**Fig. 3a**). The nine clusters produced at this threshold were enriched for proteins with relatively broad cell component annotations: nucleus, mitochondrion, Golgi apparatus, cytoplasm, endoplasmic reticulum (ER), actin patches, nucleolus and cytosolic ribosome (all p-value<10e-20 except cluster 4 with p-value<10e-5; **Fig. 3a, b;Table S1**; examples of cell images from each cluster are shown in **Fig. S2a**). At this level of feature profile similarity, proteins annotated to subcellular components with visually distinct morphologies, such as organelles, tend to be in a single cluster, whereas proteins annotated to more heterogeneous cellular compartments are found in multiple clusters. For example, proteins with a *nucleus* GO cellular component annotation are enriched only in cluster 1, whereas proteins with a *cytoplasm* annotation were enriched in clusters 4, 8, and 9 (**Fig. 3a, b**). Detailed visual image inspection revealed that some clusters reflect protein localization to both the cytoplasm and another compartment, such as cluster 4 which contains subsets of proteins localized to the cytoplasm and cell surface proteins. Other clusters likely reflect differences in protein abundance, such as cluster 8, which includes a number of highly abundant proteins, including ribosomal proteins.

Next, we derived optimal correlation thresholds on our dendrogram corresponding to two additional, more detailed biological standards: GO Slim Bioprocess and Protein Complexes (**Fig. S2b**). We obtained 21 clusters for GO Slim Bioprocesses (0.64 AMI distance threshold), 20 of which were functionally enriched (GO enrichments are shown in **Fig. S2c**; median p-value of 5e-10 across all enriched clusters; clusters’ entropies in terms of present localizations are shown in **Fig. S2d**). Similarly, 205 clusters were found at the dendrogram cutoff corresponding to a protein complex standard (0.29 AMI, **Fig. S2b**), which had 11-fold median enrichment in protein complex predictions across all clusters (distribution of the per-cluster enrichments at 0.29 AMI cutoff is shown in **Fig. S2e, f**). Hence, aFPs present robust and memory-efficient representations of protein features, which allow detection of clusters with functionally related proteins at various levels of cellular organization, with the highest functional resolution at more general levels of the hierarchy (**Fig. S2f**).

### Adaptation of PIFiA features to external annotations for protein localization and function

Our unsupervised clustering analysis showed that PIFiA aFPs capture information from cell images that enables label-free resolution of cellular spatial organization, grouping proteins by shared localization and biological function **(Fig. 3a, b)**. We have investigated another useful property of feature profiles - adaptability for subsequent supervised training. One of our goals was to create a model that produces universal feature profiles that can be used without the requirement to re-train a full neural network from scratch on a specific task. To evaluate the adaptability of feature profiles, we used the widely adopted linear evaluation protocol^16^, where a linear classifier is trained on top of the representations extracted from the network, and test accuracy is used as a measure of representation quality.

We first evaluated how PIFiA features can be adapted to protein localization labels, which is the largest labeled functional standard available^8, 10^. This analysis enables assessment of whether information contained in self-supervised PIFiA features matches the content of the original images, when extracted with a supervised method. We trained a multinomial logistic regression (LR) using PIFiA scFPs from 2415 proteins manually annotated to localize to a single subcellular localization^10^ (see Methods). We compared the final performance of the LR trained on PIFiA scFPs to DeepLoc, a supervised neural network specifically trained to classify protein localization from images of the yeast ORF-GFP collection. To match the training protocol of DeepLoc, we used scFPs derived from the single-cell images in DeepLoc’s training set^8^. We report AP scores on the same single-cell crops from the test set across the full roster of 2415 single-localized proteins (**Fig. S3a**). Remarkably, PIFiA self-supervised scFPs that were paired with LR yielded a comparable performance to the supervised network DeepLoc, even though PIFiA feature profiles are self-supervised and were fitted to localization labels solely using LR. This finding suggests that PIFiA feature profiles have rich functional content, and we can use them to predict functional protein attributes without training a full network from scratch.

To visualize adaptation of PIFiA feature profiles to the supervised localization labels, we transformed the 64-dimensional aFPs into 2-dimensional space using t-SNE^35^ and colored them according to LR localization predictions (**Fig. 3c**). Each aFP on the t-SNE projection was annotated with the localization category corresponding to the maximal LR prediction across all scFPs. In this visualization, the morphological similarity of proteins encapsulated in the aFPs was translated into proximity on the 2D t-SNE map, highlighting that the separation of self-supervised aFPs on the map was driven by subcellular localization signals. We compared the aFP localization assignments with the assignments made using supervised machine learning methods or manual annotations trained to specifically assign proteins to subcellular localizations (**Fig. S3b**). Ultimately, we observed higher quality of linear localization annotation compared to subcellular localization standards produced by other approaches^2, 8^ (**Fig S3a**).

These analyses show that PIFiA feature profiles can be adapted to the objective of a supervised neural network, which confirms the high information content of PIFiA features. Such adaptable feature profiles may accelerate training by replacing various task-specific supervised neural networks with one multi-purpose self-supervised approach, which yields universally applicable representations.

### Experimental validation of PIFiA predictions of sub-compartmental organization of the cell

We explored more specific functional information associated with PIFiA aFPs. We used Gaussian kernel density estimates^36^ (KDEs) (see Methods) to annotate our whole proteome 2D tSNE projection of aFPs with Gene Ontology bioprocess terms. For illustration, we selected terms from different subcellular components that had the lowest variance on the t-SNE map. This annotation showed that PIFiA features distinguished biological processes within cellular compartments (**Fig. 3d**). For example, regions of the tSNE map corresponding to the cytoplasm (**Fig. 3c**) had distinct regions enriched for translation initiation and elongation, P-body assembly, pentose-phosphate shunt and glycogen metabolic process (**Fig. 3d**). This analysis illustrates that GFP-tagged proteins with similar biological roles have distinguishable appearances in cell images, and that PIFiA learns feature profiles that can be used to discover protein functional groups across different levels of subcellular organization, including organelles and possible sub-compartmental structures.

To further explore information in PIFiA profiles related to the sub-compartmental organization of the cell, we clustered aFPs of proteins that mapped to the same localization category to produce 15 per-compartment hierarchical trees (derived from the 15 categories defined by LR; **Fig. 3c**). We selected a sub-compartmental clustering threshold of 0.5 based on the highest morphological similarity within clusters and maximal separation between clusters, measured by a Silhouette score (**Fig. S2e**). We identified 30 clusters, which are indicated on the tSNE projection of PIFiA feature profiles in **Fig. 3e**, with example images of cells from each group shown in **Fig. S4** (see also **Table S2** for GO enrichment and other information). We refer to these clusters using their localization category and associated group number (e.g., nuc-1 corresponds to the first sub-compartmental group in the nucleus). We provide an interactive version of the t-SNE plot from **Fig. 3e** on the PIFiA website (https://thecellvision.org/pifia/), where each point on the plot is clickable and allows the user to explore the micrographs corresponding to the GFP-tagged protein, find its nearest neighbours and perform enrichment analysis based on the closest aFPs.

Several general features associated with the clusters in **Fig. 3e** suggest that they are functionally meaningful and reflect sub-compartmental organization. First, proteins localizing to compartments which tend to be more homogeneous in their morphological patterns were typically seen in a single cluster (e.g., peroxisome, spindle pole, vacuolar membrane, nuclear periphery), while proteins associated with large or heterogeneous compartments, such as the nucleus, cytoplasm and mitochondria, defined more than one sub-compartment cluster (**Fig. 3e, Table S2a**). Second, 16 of 32 groups showed >2-fold enrichment for a GO annotation category (**Table S2b**, P<0.01). For example, the nucleus region of the whole-proteome map was divided into five clusters, enriched in GO bioprocesses such as small molecule metabolic process, chromatin remodeling and RNA polymerase II activity, mitotic nuclear division and proteolysis (**Fig 3e, Table S2**). Third, as expected, some of the groupings appeared to be based on protein features that were easily discernible. For example, *nuc*-1 clustering likely resulted from high protein abundance, and it included histones and metabolic enzymes (median GFP intensity *nuc*-1 proteins=5834±2103 vs median for all *nuc* proteins=745±793) (**Table S2**). For some of the other groups, clustering appeared to result from differences in the spatial distribution of pixels in a region. For example, *cyto*-3 proteins all had a prominent cytosolic signal overlaid with a punctate morphology, and most had roles in Golgi vesicle transport (**Table S2, Fig. S4**). Similarly, *cyto*-8 contained only seven proteins with no obvious functional overlap, but by visual inspection, all the proteins localized to the cytoplasm and to one or more foci (**Fig. S4**). Thus, a fraction of sub-compartmental groups could be explained by distinct protein localization features, which may correspond to coherent functionality.

For many sub-compartmental groups, however, the features driving the clustering were less obvious. To ask if we could manually identify differences between proteins from different PIFiA sub-compartments with the same overall localization, we used a more sensitive colocalization assay. We chose pairs of proteins with similar abundances, tagged them with two fluorescent proteins, mNeonGreen and mScarlet, imaged cells containing both tagged proteins, and manually assessed images (**Fig. 3f, Table S2d**). Using colocalization, we observed subtle differences in most of the pairs from different sub-compartmental groups; specifically, we identified differences in 39/52 (75%) protein pairs from distinct groups, but in only 7/24 (29%) pairs from the same group (**Table S2d**). For example, among nuclear sub-compartmental groups, we identified a distinct localization for *nuc*-5 proteins, which were 13.5-fold enriched for components of the proteasome (P=7.78E-22). The localization of *nuc-5* proteins overlapped extensively with that of other nuclear proteins, but *nuc-5* proteins additionally localized to the nuclear periphery (**Fig. 3f**, top row). We detected the nuclear periphery localization of *nuc-5* proteins when we looked at different proteasome components from nuc-5 in colocalization assays with proteins from different sub-compartmental *nuc* groups (**Fig. S5**).

In another example, we performed co-localization analysis with proteins assigned to different cell periphery (CP) groups. We examined cells expressing both a high-affinity iron transporter, Ftr1, from the *CP*-1 group, and Tcb2, a protein involved in ER-plasma membrane tethering, from the *CP*-2 group (**Fig. 3f**, middle row). Ftr1, tagged with mScarlet, and Tcb2, tagged with mNeonGreen, localized to distinct regions of the cell periphery. Ftr1 localized specifically to the mother cell periphery but was absent from the bud, whereas Tcb2 was present at both the mother and daughter cell peripheries. Indeed, by visual inspection, we found that all the *CP*-1 proteins had mother-specific localization, like Ftr1. In total, the *CP*-1 group contains 21 proteins, and included 7 of the 8 proteins found previously to localize asymmetrically to mother cells, all of which are members of the MDR (multidrug resistance) transporter family: Fui1, Hip1, Hnm1, Pdr5, Snq2, Tpo1, Yor1^37, 38^.

In addition to the MDR transporters, the *CP*-1 group contains 14 novel mother-specific proteins, including several other transporter proteins (Atr1, Ftr1, Hxt6, Mep1, Mep3, Qdr2, and Qdr3), proteins with roles in signaling (Gpa2, Mid2, Psr1), and three relatively uncharacterized proteins, including Ybr016w, which is a tail-anchored plasma membrane protein that is orthologous to human CYSTM1 (cysteine rich transmembrane module containing 1), Ina1, an uncharacterized protein whose paralog protein, Fat3, is required for fatty acid uptake^39^, and Crp1, an uncharacterized protein whose paralog protein, Mdg1, appears to modulate pheromone-mediated signaling and cell polarization^40^ (**Table S2b**).

In a third test, we looked at colocalization of two proteins whose ORF-GFP fusions shows some ER localization, including Ubx2, a protein from the *ER-3* cluster involved in ER-associated protein degradation^41^, and Kre1, a protein from the *ER-5* cluster that normally functions as a cell wall glycoprotein^42^. The *ER-5* protein, Kre1, shows an ER localization but it also concentrates more discretely at the cell periphery compared to Ubx2 (**Fig. 3f**, bottom row). This localization difference was observed in other members of the sub-compartmental groups, with *ER-3* proteins, which tended generally to have a more diffuse localization than proteins in *ER-5* (**Fig. S4**).

In summary, our data show that within a compartment, PIFiA features can distinguish groups of proteins with subtle differences in localization that often have different biological roles. Many of these groups are enriched for proteins that perform biological functions not previously associated with distinctive localization patterns.

### Analysis of proteins with multi-compartment localization using PIFiA single-cell feature profiles

The single cell feature profiles (scFPs) produced by the PIFiA CNN provide an opportunity to explore more nuanced protein behaviors, including proteins localizing across multiple compartments. Previous analyses of the yeast ORF-GFP collection showed that a large fraction of the proteome localizes to two or more compartments in the same cell^2, 8, 10^. These studies used average statistics for cell populations, precluding differentiation of proteins that localize to multiple compartments, or those that shuttle between compartments. We annotated scFPs of every protein with localization categories using LR classification scores, and then we investigated the distribution of each protein’s single-cell localization scores, focusing on the two most probable localizations (see Methods). Using this strategy, we found that most (3424) proteins have a homogeneous localization (localizing to a single compartment), while 652 proteins exhibit localization heterogeneity (localizing to two or more compartments) (**Fig. 4a**). We classified the proteins with heterogeneous localization into two categories: (1) 396 proteins that localize to more than one compartment in a single cell, which we refer to as AND-proteins, and (2) 256 proteins that appear either in a primary or a secondary location but not in the same cell, which we refer to as OR-proteins (**Fig. 4b, c, d Table S3**). For most proteins the assigned localization probabilities are continuously distributed, but our classification summarizes the localization, indicating the compartments that the protein predominantly populates. For example, Pho85, is classified as an AND-protein with a mixed signal predominantly from nucleus and cytoplasm within single cells, consistent with its known biology^43^ (**Fig. 4c)**. In contrast, Stb1 is a transcription factor whose nuclear localization is cell cycle regulated and it was classified by our analysis of scFPs as being either nuclear or cytoplasmic (OR-protein), as seen in previous studies^44^ (**Fig. 4c**).

**Figure 4.**
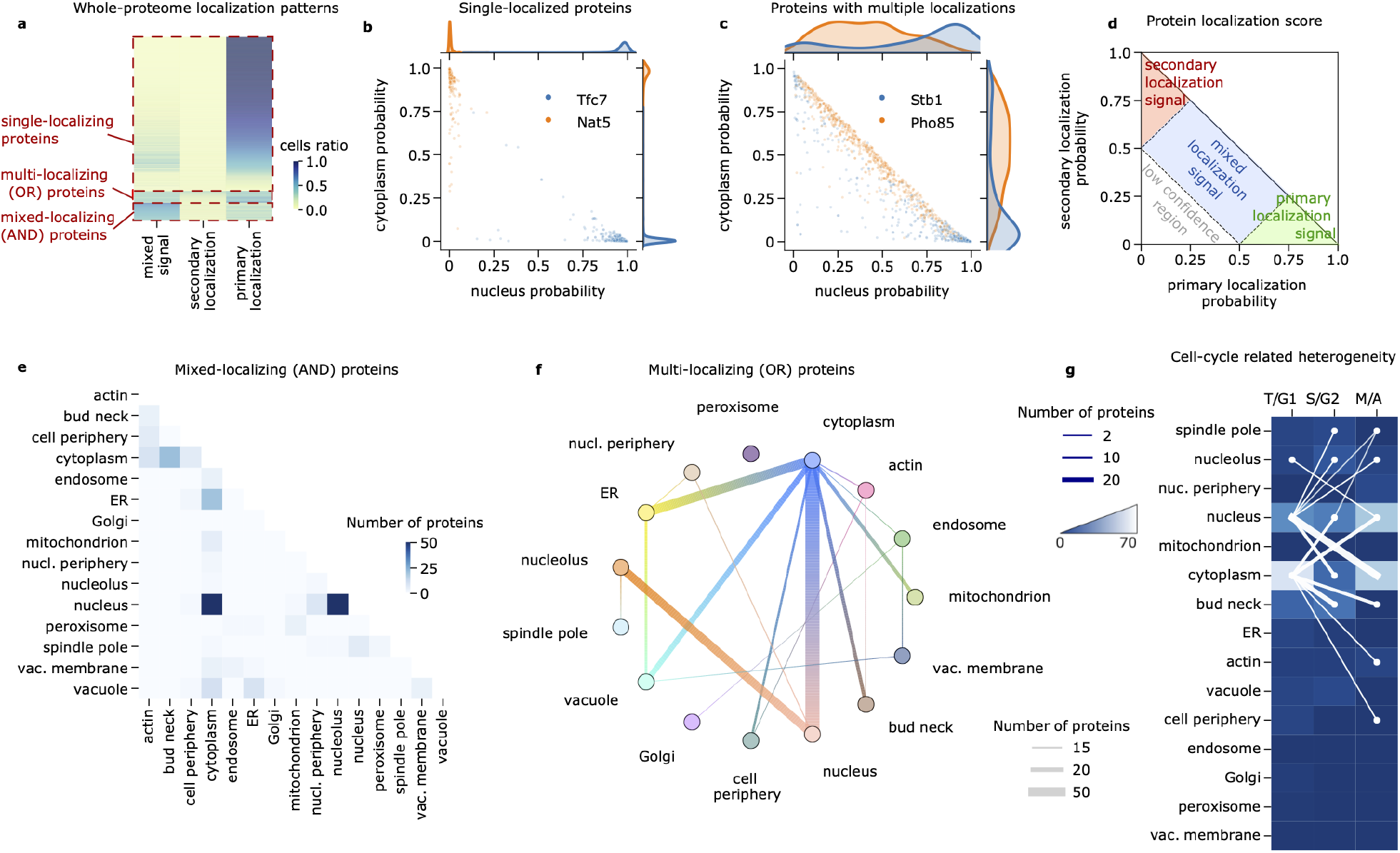
Identification of proteins with morphological heterogeneity using PIFiA single-cell feature profiles. **a**, Summary of whole-proteome analysis of localization heterogeneity. The heatmap depicts ratios of cells falling into a mixed localization category, secondary and primary localization regions (following the introduced scoring schema). Proteins are arranged in the following order top-to-bottom - proteins with homogeneous localizations, OR-type proteins and AND-type proteins. Proteins with low ratios in all three columns (yellow color) feature many cells that fall into the low confidence region. **b**, Dissecting localization heterogeneity of a protein at a single-cell level: homogeneous localization. Examples of two proteins with homogeneous localization patterns are illustrated using a scatter plot. Each point on the scatter plot corresponds to a single-cell crop, mapped to the probability of nuclear and cytoplasmic localization according to the LR predictions. In this example, the majority of Nat5-GFP expressing cells exhibit clear cytoplasmic localization, while most Tfc7-GFP expressing cells show a clear nuclear pattern. **c**, Dissecting localization heterogeneity of a protein at a single-cell level: heterogeneous localization. Examples of a protein with AND-type localization heterogeneity (Pho85, single-cell FPs shows patterns from its primary AND secondary localization), and multi-localizing heterogeneity (Stb1, protein can be in either its primary OR secondary localization, but not both at the same time) are illustrated using a scatterplot. Cells expressing the Pho85 protein show a mixed signal from the nucleus and cytoplasm, with most cells being distributed along the diagonal. In contrast, Stb1 has two populations of cells in the scatter plot centered either around the nuclear or the cytoplasmic corner. **d**, Schema of scoring proteins for localization heterogeneity using a single-cell level distribution of localization probabilities. Probabilities are obtained from primary and secondary localizations (i.e. first and second most probable localizations of the logistic regression classification of that protein). **e**, Localization co-occurrence heatmap for 396 AND-localizing proteins, showing numbers of proteins present at two localizations. The scale bar is set to a maximum intensity of 50 to enable visualization of categories with fewer proteins (see quantities in **Table S3**). **f**, Circle plot depicting localization patterns of 256 OR-type proteins. Thickness of the line connecting two localizations indicates the number of proteins showing localization heterogeneity between these localizations. **g**, Localization heterogeneity related to cell cycle position for 136 proteins, which exhibited statistically significant cell cycle variation. Connections indicate localizations of the proteins that are present at specific cell cycle stages (thicker lines indicate a more common connection between particular localization change and cell cycle stage transition). The color of the heatmap indicates the number of heterogeneous proteins that are present in the corresponding cell cycle phase for a particular localization.

We summarized overall AND-/OR-localizations across 15 localization categories, which clearly illustrated that a large fraction of these changes involved the nucleus and cytoplasm compartments. Among the 922 proteins with a high confidence nuclear localization, 708 were solely nuclear, 159 nuclear AND cytoplasmic, and 55 nuclear OR cytoplasmic (**Fig. 4e, f**). We asked how these classes were distributed in different bioprocesses involving the nucleus^31^. As expected, proteins with roles in RNA processing and chromatin organization tended to be solely nuclear (**Table S3**). The trends for proteins that have dual localization were also consistent with well-established biology. For example, proteins with roles in cell cycle progression were 4.1-fold enriched in nucleus OR cytoplasm (P=6.7E-05). Many cell cycle proteins, in particular many transcription factors, use localization to regulate protein activity^46^. Proteins with roles in DNA replication/repair and stress response were weakly enriched in nucleus AND cytoplasm (1.5-fold, P= 1.40E-03 and 1.9-fold, P=9.7E-04 respectively). DNA repair proteins often alter their relative localization in the presence of damage, either to initiate a repair response or to prevent catastrophic cell cycle events^47^. Because our cells were not experiencing DNA damage at the time of imaging, many of these proteins displayed both nuclear AND cytoplasmic localization in our data. Hence, while many protein groups that show different patterns were too small to perform consistent enrichment analysis, enrichments that were seen for nuclear-cytoplasmic groups, where there are enough proteins to assess, were consistent with known biology.

Finally, because proteins with roles in cell cycle progression were enriched among both the OR- and the AND-proteins, we used our scFPs to assess how cell cycle position could account for some of the protein localization heterogeneity. To do so, we took advantage of the nuclear and cytoplasmic markers (td-Tomato-NLS; E2-Crimson) in our GFP collection to explore the relationship between cell cycle position and protein abundance or localization heterogeneity. We first trained an ensemble of CNNs on the nuclear and cytoplasmic RFP channels to predict one of four cell cycle stages, and we subsequently mapped each single-cell crop to a cell cycle category: T/G1 (Telophase, Gap phase 1), S/G2 (DNA synthesis phase/Gap phase 2) or M/A (metaphase/anaphase) (see Methods). We then compared the single-cell distributions of each cell cycle stage with the localization calls to discover relationships between protein localization and cell cycle (**Table S3**). In total, we identified 136 proteins with cell-cycle-dependent variation in PIFiA feature profiles, determined by Mann-Whitney U test^48^ (p-value<1e-3, see Methods). Our results are summarized in the connected heatmap shown in **Fig. 4g**. As expected, some of the discovered localization changes reflected cell-cycle-dependent differences in the corresponding compartment. For example, most bud neck/cytoplasm AND-localizing proteins (14/23 proteins) were cytosolic in G1 before the bud neck had formed and localized to the bud neck later in the cell cycle (**Table S3**). However, many cell cycle regulated proteins moved between permanent compartments, including 66 moving between the nucleus and cytoplasm. Indeed, PIFiA identified 4 proteins not previously known to be cell cycle regulated, that localized to the cytoplasm and nucleus, but showed a predominantly cytoplasmic localization in M/A (Yel025c, Atc1, Bop3, and Cmg1).

Overall, scFPs derived from the self-supervised PIFiA workflow enable resolution of single-cell localization and are suitable for cell-to-cell variability analysis. Notably, PIFiA feature profiles contain enough functional information to distinguish compartments and sub-compartmental morphologies without pre-assigned labels, enabling analysis of protein localization heterogeneity in a data-driven way, which precludes propagating annotation errors.

### Prediction of functional modules using PIFiA single-cell feature profiles

AMI scores at different correlation thresholds allowed us to resolve functional information associated with hierarchical clustering of PIFiA aFPs (**Fig. 2a**) and determine an optimal correlation threshold for discovering functional modules, such as protein complexes (**Fig. S2a**). However, averaging feature profiles leads to information loss, which is not optimal for more precise analysis. Hence, we explored the use of single-cell feature profiles for the identification of functional modules. In particular, we focused on protein complexes, which represent functional modules whose components are expected to colocalize within a single cell.

To visually investigate protein complex distributions with scFPs, we projected scFPs from the test set using 2D tSNE (**Fig. 5a**). The scFPs of proteins from the same complex often localized together on the tSNE map, but they were typically intermingled and difficult to separate from each other, which is consistent with the resolution of light microscopy. Nevertheless, the scFPs corresponding to different protein complexes with the same subcellular localization were often separated on the tSNE map (e.g., Polymerase-II, Polymerase-III and RSC in the nucleus; EGO and V-ATPase in the vacuolar membrane) (**Fig. 5a**).

**Figure 5.**
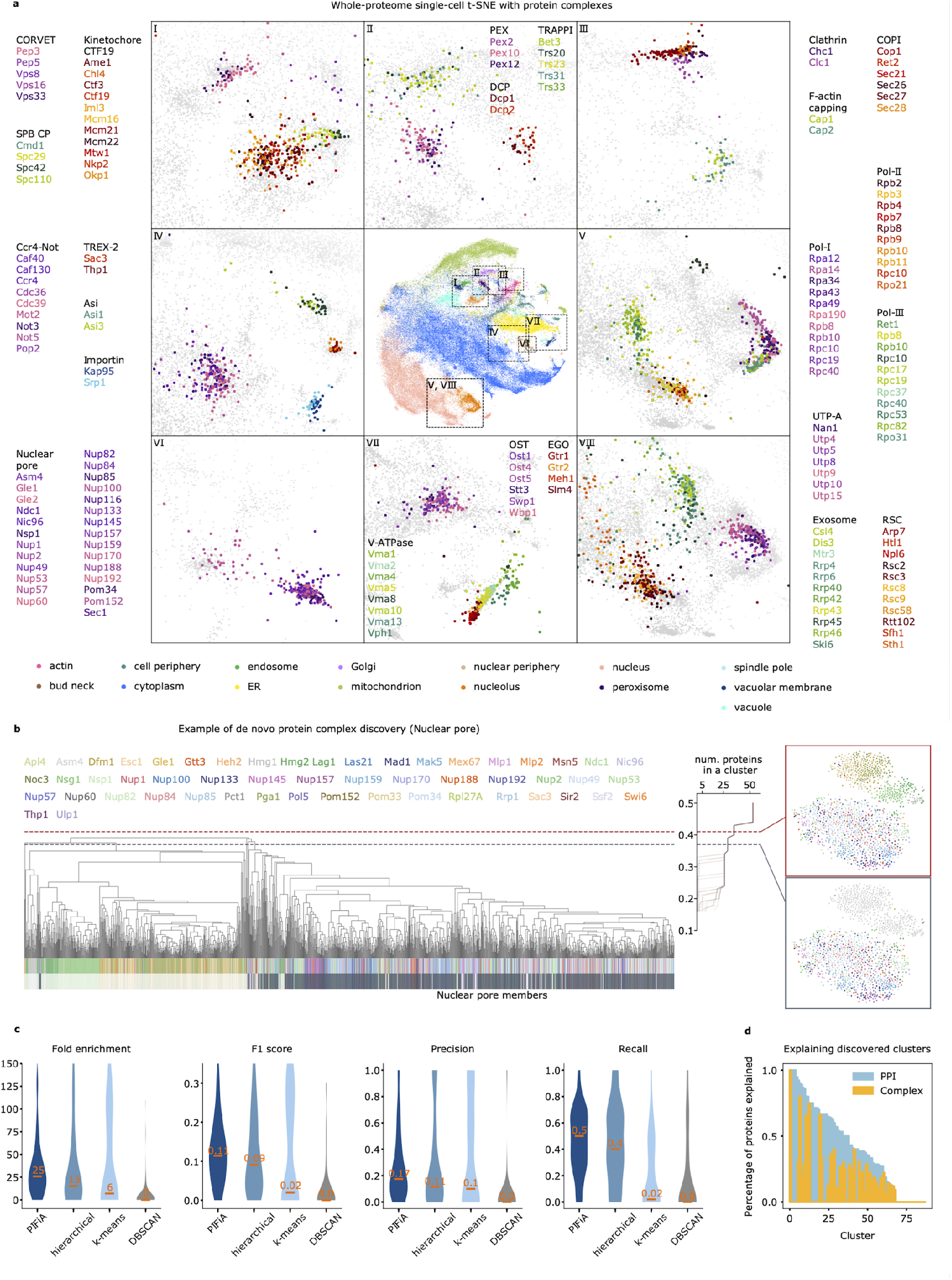
Discovery of protein functional modules using PIFiA single-cell feature profiles. **a**, Visualization of protein complex clusters on a single-cell tSNE plot of PIFiA feature profiles. The central plot shows a whole proteome tSNE projection of PIFiA single-cell feature profiles (scFPs). Each point on the plot represents a protein that is colored according to 15 different subcellular localizations (color codes are explained below the plot). Zoom-in plots show a more detailed view of some regions of the global tSNE, showing single-cell features from the test set corresponding to proteins from the same complex with the same color palette, each protein shown in different color. **b**, The scFPs dendrogram shows clustering of scFPs highlighting a region that identifies a nuclear pore cluster among nuclear periphery scFPs.The line graph on the right shows changes in the number of proteins in a cluster (X axis) when the correlation threshold for clustering is changed (Y axis). Zoom-in plots of two clusters at different correlation thresholds (red and grey dashed lines) are shown as scFPs tSNE plots to the right. **c**, Violin plots comparing the performance of four clustering approaches on 140 protein complexes are shown: our clustering using PIFiA scFPs, hierarchical clustering, k-means and DBSCAN. **d**, Plot illustrating the fraction of proteins in each cluster with a protein-protein interaction annotated in the BioGrid (blue) or as protein complex member (yellow). Percentage of cluster proteins participating in a protein-protein interaction or being part of the protein complex are shown on the y-axis. Cluster numbers are sorted based on the number of discovered interactions.

To further explore the utility of scFPs for algorithmically identifying protein complexes, we developed a modified hierarchical clustering approach, called adaptive thresholding, that is designed to identify correlation thresholds on the hierarchical dendogram at which scFPs inside a cluster become indistinguishable, and thus might be expected to contain interacting proteins (see Methods). We performed hierarchical clustering of test set scFPs using average linkage and a correlation metric. We then traced the number of unique proteins inside the cluster along with the divisions of the single-cell dendrogram to discover levels of the dendrogram at which the number of proteins in a cluster plateaus (**Fig. 5b**). Such plateaus identify levels of the global scFPs dendrogram at which single-cell features are practically inseparable, a division threshold that we call a root cluster (see Methods). For example, our adaptive thresholding method identified a root cluster corresponding to the nuclear pore complex, which distinguished it from other nuclear periphery proteins **(Fig 5b)**.

We compared the adaptive thresholding method to other clustering approaches from three different families - connectivity (hierarchical clustering^33^), centroid (k-means^49^) and density methods (DBSCAN^50^) (**Fig. 5c**). For each of the methods used in our comparison, we tried a range of hyperparameters and selected the ones that maximized median F1-score (see Methods). Evaluation was performed on a set of 140 protein complexes that contain at least three proteins included in the ORF-GFP localization dataset^31^. Our approach outperformed other methods in terms of four different scores - fold enrichment, F1 score, precision, and recall (**Fig. 5c**). The distributions of scores highlight the advantages of our adaptive thresholding approach. Density-based clustering fails at the protein complex identification task due to the high density of the feature space. At the same time, k-means fails at the identification of larger protein complexes (more than 5 protein members), hence its violin plot has two peaks. Hierarchical clustering is a more advantageous approach for this task, yet it requires information on the distance threshold and lacks adaptability for the protein complex size and cellular compartment. In contrast, our adaptive thresholding approach finds an optimal distance threshold for each cluster and, hence, it can discover protein complexes of varying sizes.

Using the adaptive thresholding clustering approach, we constructed a list of 88 high-confidence clusters whose proteins were indistinguishable at the single cell level (**Table S4, Fig. 5d**). We mapped each cluster to a protein complex with the maximal fold enrichment and saw a median fold enrichment of 36.5 across 88 clusters, which is a 3-fold improvement over our clustering of aFPs with an optimized cut-off (**Fig. 2**). Of the 88 predicted clusters, 43 captured members of 32 different protein complexes distributed across 15 subcellular compartments (**Table S4**). The remaining clusters did not capture two or more members of the same protein complex, although in 25/45 cases they contained PPIs (as annotated in BioGrid^51^). By using proteins from the same localization as our background set to compute fold enrichment, we tested whether the clusters could differentiate a protein complex from others in the same subcellular location (see Methods). We discovered that PIFiA confidently predicted members of protein complexes in multiple compartments, such as: [1] the proteasome and Ada2/Gcn5/Ada3 transcription activation complex in the nucleus; [2] LSM2-7 complex, decapping complex and translation initiation factor 2B complex in the cytoplasm; [3] the oligosaccharyl transferase and Sec62-Sec63 complexes in the ER; [4] vacuolar proton translocating ATPase complex, phosphatidylinositol 3-kinase complex and iron exporter complex in the vacuolar membrane; [5] F-actin capping protein complex and PAN1 actin cytoskeleton-regulatory complex in the actin cytoskeleton; [6] Spc105 complex and NDC80 complex in the spindle pole; [7] retromer complex and SNX4-SNX41 sorting nexin complex in endosomes (see **Table S4**).

In some cases (17/44), we identified all members of a complex, together with some additional proteins, which may be previously unappreciated complex components or members of an extended functional module, such as regulators or target proteins. For example, cluster #6 contained all 4 subunits of the COMA complex, a kinetochore component that connects proteins bound to centromeric DNA with those bound to microtubules, as well as nine additional proteins, eight of which display protein-protein interactions (PPIs) with COMA members^51^. Other clusters identified proteins with the same biological role, that may participate in PPIs. For example, cluster #39 contained 26 proteins that localized to the nuclear periphery in a punctate fashion. This group contained the only two GFP-tagged members of the TREX-2 complex, which couples SAGA-dependent gene expression and transcription elongation to mRNA export at the nuclear pore complex. Cluster #39 also included 8 proteins reported to have PPIs with members of the TREX complex, suggesting they may function in concert with the complex^51^. Among the remaining proteins were members of a silencing complex, including Sir2, Sir3, Sir4 and the Sir4-interacting protein Esc1, which suppresses transcription at subtelomeric regions, tethering them to the nuclear periphery^52^. Thus cluster #39 identified proteins with roles in gene expression that localize to the nuclear periphery, some of which function to modulate each other’s activity.

In summary, we have developed a novel approach for discovery of functional modules using solely the self-supervised feature profiles and leveraging the properties of microscopy data for optimal clustering, and prediction of molecular interactions.

### Interpretation of PIFiA features

Our analysis shows that PIFiA feature profiles contain condensed information about protein function at various levels of granularity. However, since deep neural networks function as ‘black-box’ models, it is difficult to dissect feature profiles and explain how individual features are related to the input images. To attempt to interpret PIFiA features, we first quantified feature importance for 15 different localization categories covering a diverse set of subcellular morphological patterns. We used the LR model described earlier to derive importance scores for each feature; the coefficients of the trained LR quantify how much each feature is predictive of a certain localization (**Fig 6a**). Most localizations had more than three strongly predictive features (LR coefficient value >5), suggesting that PIFiA learns to detect several distinctive patterns for each subcellular compartment. This confirms that PIFiA learns localization patterns with its convolutional filters, despite being trained on a completely different self-supervised objective. Also, the same feature could be predictive for several localizations (for example, features #3 and #28 recognize circular patterns corresponding to the vacuolar membrane and nuclear periphery), or react to some variation of visual patterns present in multiple localizations. Overall, larger and more complex compartments required more features to be confidently classified. To illustrate this finding, we plotted classification accuracy for the three largest compartments (cytoplasm, nucleus and mitochondria), as well as three homogeneous compartments (nucleolus, peroxisome and vacuolar membrane) with respect to the number of features used during LR training (see Methods) (**Fig. 6b, c, S7a**). While larger localization categories required approximately 30 features to reach their best performance, smaller localizations reached a saturation point at around 10 features.

**Figure 6.**
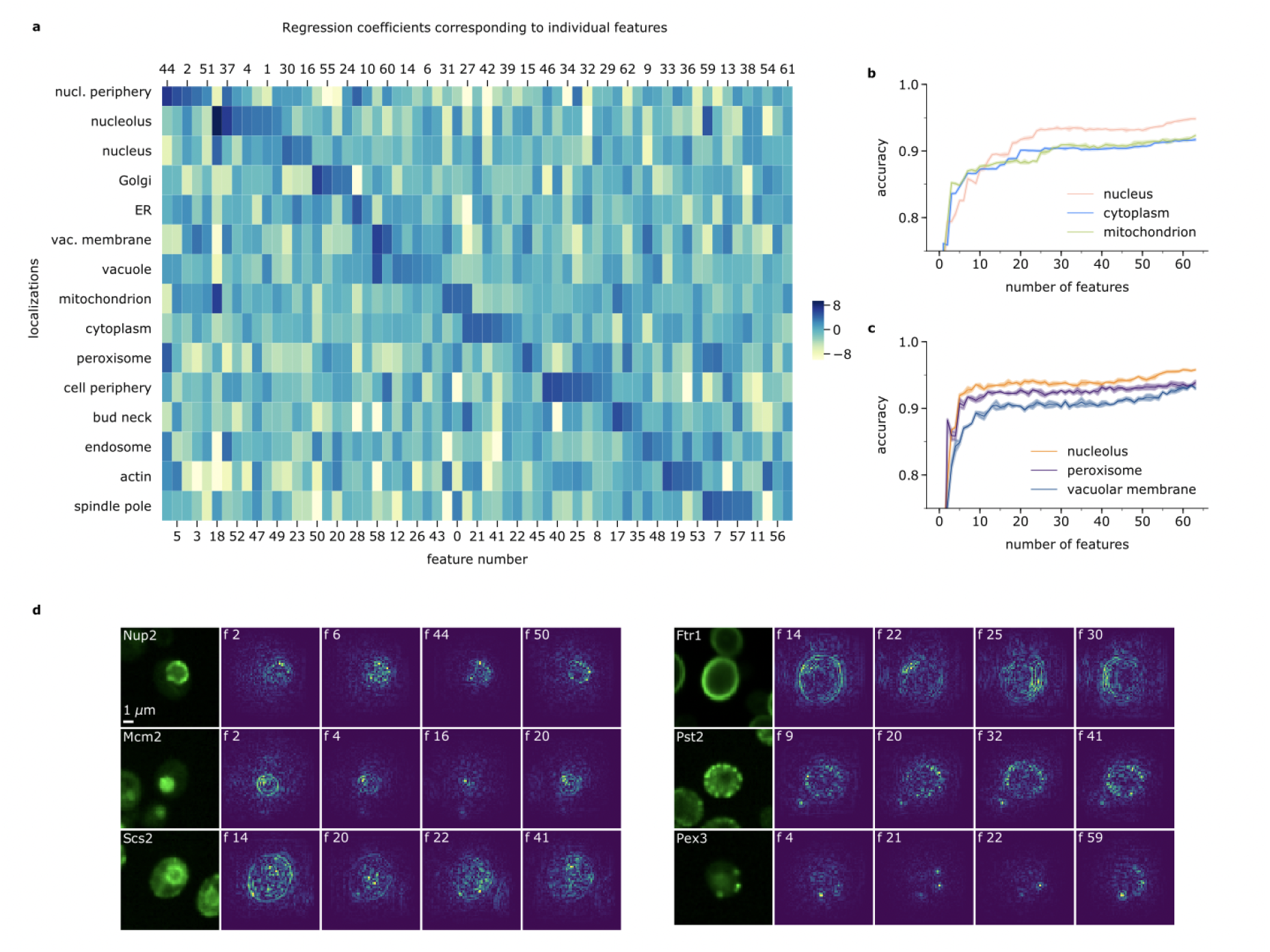
Interpretation of PIFiA features. **a**, Heatmap of the logistic regression coefficients for localization prediction associated with each feature. Each feature was mapped to a localization (based on the maximal coefficient value), and features were sorted in descending order of coefficients. **b, c**, Classification accuracy depends on the number of features used to train the logistic regression. We trained logistic regressions with varying numbers of features in the representation profile (sorted in the order of importance for the localization). The three largest and most complex localizations are shown in b, while three smaller and more homogeneous localizations are depicted in c. Shading shows standard deviation across 5 runs. **d**, Gradient maps highlight regions of the input image that CNN “pays attention to”. We show proteins from six distinct subcellular compartments and their gradient maps obtained from four different features (most visually distinct gradient maps with the highest activation values are shown). Features result in different gradient signals, confirming that the CNN captures a variety of within-cell morphological patterns.

Another way to interpret features learned by a CNN is to find regions of the image that had a large influence on the final result^53, 54^. Using gradient calculations, importance scores can be assigned to the input image pixels depending on the degree to which they affect the classification result or individual feature values (see Methods). We used the SmoothGrad^54, 55^ approach to construct gradient maps for several features of the same protein image. We selected proteins representing five distinct subcellular localizations: Nup2 from nuclear periphery, Mcm2 from nucleus, Scs2 from ER, Ftr1 and Pst2 from cell periphery, and Pex3 from peroxisome. For each of the proteins, we used its scFPs to calculate 64 gradient maps for each of the individual features and selected four visually distinct gradient maps with the highest activation values for illustration purposes (**Fig. 6d, S7b**). We observed that different features of the same image resulted in different gradient maps. For example, gradient maps of the Mcm2 protein highlighted the nuclear periphery region, focus points in the nucleus and nuclear background signal. Similarly, different features of Ftr1 reacted to various subregions of the cell periphery. Of note, the generated gradient maps showed that the region of network attention was always the single central cell of the crop even for crops containing more than one cell, confirming that per-crop feature profiles are in fact single-cell profiles (**Fig. 6d**, with Mcm2, Ftr1 and Pst2 proteins containing multiple cells in their crop, **S7b**). Gradient-based interpretability approaches are useful to explain the relationship between individual features in the feature profile vector and input pixels in the image, and they constitute an important component of our downstream analysis pipeline. Hence, despite PIFiA’s self-supervised training objective, we can visually understand what each learned feature represents in terms of the input image regions.

## Discussion

We describe PIFiA, a self-supervised computational workflow that learns protein functional signatures from single-cell fluorescence microscopy data. Feature profiles learned by PIFiA show state-of-the-art performance on a variety of biological functional benchmarks, outperforming existing approaches for protein representation learning. Notably, our approach does not require any labels or annotations during training and uses only a single fluorescent channel. Hence, PIFiA can be easily applied to virtually any imaging dataset. We pre-trained PIFiA on a large-scale dataset encompassing over three million single-cell images of yeast cells expressing 4,049 GFP-fusion proteins – a scale comparable to that of the commonly used computer vision dataset, ImageNet^56^. As with ImageNet, we show that the yeast ORF-GFP dataset is a source for high-quality representation learning, enabling PIFiA to learn universal feature profiles that can be used out-of-the-box or minimally fine-tuned to suit an external standard. Thus, PIFiA can accelerate the rate of supervised training on external tasks by producing feature profiles that can fit any downstream task with simple linear regression, replacing multiple task-specific convolutional networks.

The PIFiA workflow unites a self-supervised convolutional neural network with multiple techniques for downstream feature profile analysis. The key advantage of our self-supervised objective is its independence of human annotations and its ability to learn high-quality features and ignore imaging artifacts and cell positions^15, 22, 57^. To ensure that PIFiA remembers solely the biologically relevant patterns, yet ignores cell positioning and replicate noise, we require the network to learn the actual GFP-tagged protein by predicting its identify. Overall, we show that features learned by PIFiA outperform another self-supervised method, Paired Cell Inpainting^15^, that was used to analyze the yeast GFP collection and even reaches the performance of the supervised approach, DeepLoc^8^, in its target task.

We describe downstream analysis techniques that use PIFiA feature profiles to explore different levels of subcellular organization that span both protein-level and single-cell feature profiles. Of note, our analysis focuses not only on the construction of a whole-proteome hierarchical map, but also provides quantitative rules to obtain clusters corresponding to a specific level of cellular organization, such as sub-cellular localization or biological process, and to identify proteins with multiple localizations and interacting proteins. This type of unbiased analysis can reveal unexpected properties and potential functions of proteins that can be further explored experimentally. For example, we used PIFiA features taken from images of yeast cells expressing GFP-tagged proteins to identify sub-compartmental groups enriched for proteins with biological processes not previously known to have distinctive subcellular localization patterns (**Fig 3e, f**). For example, we found that in addition to the known pan-nuclear localization, proteasome components are also localized at the nuclear periphery, a result we confirmed with co-localization experiments (**Fig. S5**). Nuclear periphery localization of proteasomes has not been reported in yeast, but an in situ cryo-electron tomography study in *Chlamydomonas* found nuclear 26S proteasomes crowding around nuclear pore complexes^58^. The role of proteasomes at the nuclear periphery may be to regulate transcription and/or to degrade proteins transiting the nuclear pore complex^58^.

We also identified a group of proteins that localized specifically at the cell periphery of mother cells and were depleted from the growing bud. Budding yeast divide asymmetrically, with a replicative lifespan of 20-30 generations, where each division gives rise to a daughter whose replicative lifespan is reset and a mother who continues to age^59^. Mother-specific cell periphery localization is achieved when three conditions are fulfilled^37^. First, mother-specific proteins lack diffusive mobility in the plasma membrane. Second, newly synthesized proteins are deposited specifically in the growing bud. Third, the genes encoding these mother-specific proteins are expressed late in the cell cycle, so for cells in S/G_2_ (small-budded cells) protein is detectable only in the mother. These steps ensure that the new and old pools of these proteins become spatially segregated during asymmetric division. Indeed mother-specific localization of cell periphery proteins has been proposed to play a role in aging, with the daughter cell getting the newly synthesized copies of the protein, and the older and potentially more damaged copies inherited by the aging mother^37^. Our set of asymmetrically segregating proteins includes 7 proteins previously seen to have mother-specific localization, plus 14 novel mother-specific proteins, including other transporters, proteins with roles in signaling, and 3 uncharacterized cell surface proteins (**Table S2**). It is possible that accumulation of old and damaged versions of these newly identified proteins may also play a role in mother-specific aging. We also applied PIFiA features for the identification of interacting proteins and members of protein complexes. To accomplish this, we used an adaptive thresholding method for single-cell clustering that exploits the biological properties of protein-protein interactions and microscopy data, outperforming conventional clustering methods for identifying members of protein complexes. We show that proteins whose single-cell PIFiA features are indistinguishable can be members of the same protein complex, have PPIs with each other, or have functionally related biological roles. A similar approach, *cytoself*, was used to visually separate protein complexes from different compartments in human cells on tSNE. We demonstrate that PIFiA can distinguish protein complexes from the same compartment in yeast cells, which are 5 to 30-fold smaller in size, providing a quantitative approach for downstream analysis and identification of functionally related proteins.

In summary, the PiFiA pipeline extracts high quality functional information about proteins from cell images in a quantitative form, without relying on a pre-existing labels or manual annotations. In essence, the approach performs *in silico* colocalization, and it can be used to identify novel properties of proteins, including similarity and variability, that have the potential to inspire new experiments to uncover novel biological insights.

## Methods

### Construction of mutant arrays for Imaging

For imaging screens, BY5299 (*MAT*α *his3Δ1 leu2Δ0 ura3Δ0 met15Δ0* lyp1pr::TDH3pr-E2-Crimson::HPH::*lyp1*Δ *can1*pr::TDH3pr-tdTomato-NLS::*URA3*::*can1*Δ::STE2pr-*LEU2)* was used as the starting query strain. E2-Crimson and td-Tomato-NLS are used as cytosolic and nuclear markers respectively. The starting strain was crossed to the *MAT***a** ORF-GFP collection^10^ and haploid strains carrying both the red fluorescent protein markers and the ORF-GFP were selected using the SGA method^28^. All SGA selection steps were conducted at 30°C, except sporulation, which was conducted at 22°C for 10 days. The screen was performed in two biological replicates.

### High-throughput microscopy

Yeast cultures were prepared for microscopy and imaged as previously described^2, 3, 6, 60^. Briefly, haploid wild-type *MAT***a** strains expressing fluorescent protein fusions from SGA final selection plates were grown at 30°C in low fluorescence synthetic minimal medium with Geneticin (200μg/mL) and Noursoethricin (100μg/ml). Cells were transferred to 384-well PerkinElmer CellCarrier Ultra imaging plates and centrifuged for 1 minute at 500 g before imaging. Micrographs were obtained on an Opera Phenix (PerkinElmer) automated spinning disc confocal microscope. All imaging was done with a 63× water immersion objective. GFP was excited using a 488 nm laser and emission collected through a 520/35 nm filter. tdTomato was excited using a 561 nm laser, and emission collected through a 600/40 nm filter. E2Crimson was excited using a 640 nm laser, and emission collected through a 690/50 nm filter.

### Image acquisition for co-localization experiments

Protein pairs were chosen for co-localization if they had similar abundance^27^ and localized to the same general subcellular compartment. For each protein, C-terminal fusions to both mNeonGreen and mScarlet were constructed as previously described^61^. Haploid cells in both configurations were mated to construct **a**/α diploids containing proteins tagged with the two fluorescent proteins. Diploid cells were grown and imaged in low fluorescence synthetic minimal media^62^ supplemented with Hygromycin B (300mg/mL), Geneticin (300mg/mL) and 2% glucose. Cells were grown at 30°C to mid-logarithmic phase and transferred to Concanavalin A-coated 384-well PerkinElmer CellCarrier Ultra imaging plates. Images were acquired at 22°C using the Opera Phenix (PerkinElmer) automated spinning disc confocal microscope. Three image fields of 5 Z-stacks of optical sections 0.7μm apart were taken for each well. Each field contained 100-150 cells, acquired using the 63x water immersion objective. mNeonGreen was excited using the 488 nM laser, with emissions collected through a 520/35 nm filter. mScarlet-I was excited using the 561 nm laser, with emissions collected through a 600/40 nm filter. Digital Phase Contrast was used for cell detection using LED bright field imaging. All images were assessed by visual inspection.

### Dataset overview and image preprocessing

Images of the 4049 strains expressing a GFP-tagged protein visible above background fluorescence were obtained using an automated confocal microscope as described above. Cell images were obtained from two biological replicates, each of which had four fields of view for each GFP-tagged strain.

As the first step of preprocessing, we computed cell centers’ coordinates across all images in the dataset using the nuclear channel. We obtained coordinates of the cells’ centers by segmenting the nuclear channel with a simple Watershed algorithm and computing x, y coordinates of the center of each cell’s nucleus^7^. We ignored cells with centers too close to the crop’s boundary (less than 10 pixels). Based on the cell center coordinates, we created single-cell crops of 64×64 pixels around those centers across all images in the dataset. We filtered crops that had GFP signal intensity less than the 5th percentile of the whole-proteome GFP intensity distribution, and crops dominated by the background noise (i.e., a uniform signal across the whole crop, with variance). After filtering low-quality crops, we dropped proteins with less than 10 cells, and we obtained 3,058,961 unique cells in the dataset. Then, the dataset was split into training, validation and test sets using 80%, 10% and 10% of the cells of each protein respectively. The training subset contained 2,450,801 single-cell crops, and validation and test subsets contained 304,080 single-cell crops each. Finally, we applied instance normalization by standardizing the raw pixel intensities of every crop to a mean of 0 and a variance of 1 (independently for each channel of each sample). PIFiA was trained on 64×64 pixel crops of the GFP channel. During training, we used random flipping (horizontal and vertical) and random rotation across {0, 90, 180, 270} degrees to augment the training data. Labels of the training set are one-hot class vectors of length 4049.

### Architecture and training

The architecture details are illustrated in **Fig. S1a**. The backbone of PIFiA consists of eight convolutional blocks followed by three fully-connected layers. Each convolutional block consists of a convolutional layer, batch normalization and rectified linear unit activation. Training was performed using Adam optimizer^63^ with a learning rate of 1e-3 and cosine decay learning rate schedule (number of steps equal to the number of training updates during 30 epochs), with cross-entropy as an objective function (*y*_*i*_ and 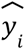 are predicted probability and ground truth label of the protein i; N is the total number of classes, i.e.,4049 proteins):

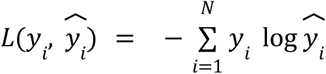

To prevent overfitting, we applied dropout regularization^64^ of 0.05 (5% dropout rate) after the second fully-connected layer (feature extraction layer). We performed hyperparameter optimization and selected the learning rate from {1e-4, 3e-4, 5e-4, 8e-4, 1e-3, 3e-3, 5e-3} and dropout rate from {0.01, 0.02, 0.05, 0.1, 0.2, 0.3, 0.5} based on maximal validation accuracy. Network parameters were initialized using a truncated normal distribution function with a standard deviation of 0.1. To report the performance, we ran the model three times with different random weights initializations; each run was 30 epochs and model weights were saved after every epoch. All the experiments were performed in Python using Tensorflow. The model was trained on the computing cluster of the Vector Institute for Artificial Intelligence, using NVIDIA T4 GPU with 12GB of VRAM, and up to 32GB of system RAM (single CPU). Source code and usage examples are available at https://github.com/arazd/pifia.

We used early stopping to select the final model^65^. We defined stopping criteria based on the model’s test accuracy of proteins classification across 4049 protein classes. We stop at an epoch where a derivative of the test accuracy becomes smaller than a threshold of 0.5% for at least 3 epochs, i.e., a point at which accuracy starts to saturate (**Fig. S1b**). Our goal was to stop at the point when the model has already grasped the most important morphological patterns, yet highly related and interacting proteins are not distinguished from each other. This trend is further illustrated by plots of average precision, F-score and precision (we show 0.9 threshold) for protein complexes and pathways standards (**Fig. S1b**). With protein prediction accuracy increasing over the course of training, the precision improved, but after some epochs, AP and F-score either saturated or started to decline. We found that accuracy saturation thresholds between 0.2% and 0.7% yielded comparable and optimal solutions, though other stopping points can be used depending on the training schedule, as well as model applications and goals. The proposed early stopping strategy helped to prevent memorizing noise and unnecessary patterns, while retaining morphologically similar proteins close in the feature space.

### Benchmarking and baseline feature extraction

We compared performance of feature profiles learned by PIFiA to features from three other popular methods for protein representation learning / extraction - DeepLoc^8^, Paired Cell Inpainting^15^ and CellProfiler^7^.

A classic modular feature extraction tool, Cell Profiler, was applied to the GFP and cytoplasmic channels of the test images across 4,049 GFP-tagged proteins. We obtained 433 pre-defined CellProfiler features that quantitatively measure cellular phenotypes, including intensity, shape and texture. Since some of the CellProfiler features can be repetitive, its representations are often post-processed with Principal Component Analysis^66^ (PCA). In our work, we evaluated both the original CellProfiler representation with 433 individual features, and its PCA projection (37 individual features) that explains 99% of the variance.

We used the DeepLoc model by Kraus et. al.^8^ as our supervised learning baseline. We trained DeepLoc from scratch on the GFP channel of the same training set images using 1,432,774 single-cell crops from 15 one-hot localization categories derived from Huh et. al.^10^ We performed hyperparameter optimization and selected the most optimal learning rate from {1e-4, 3e-4, 5e-4, 8e-4, 1e-3, 3e-3, 5e-3} and dropout rate from {0.01, 0.02, 0.05, 0.1, 0.2, 0.3, 0.5}. We chose 3e-4 learning rate with cosine decay learning rate schedule and 0.05 dropout rate based on maximal validation accuracy. The model was trained with Adam optimizer for 30 epochs (model weights were saved every epoch for subsequent evaluation), with cross-entropy as an objective function, *y*_*i*_, 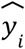 are predicted probability and ground truth label of localization i, total N=15 localization classes):

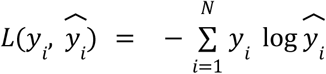

We also used early stopping to select the final DeepLoc model weights. DeepLoc model selection was based on maximal validation set accuracy. Network parameters were initialized using a truncated normal distribution function with a standard deviation of 0.1. We performed 3 runs with different random weights initializations and performed training with a batch size of 128. After training, we extracted features of the test set images from the last hidden layer of the DeepLoc model following previous studies^15, 67^.

For our self-supervised learning baseline, we used the Paired Cell Inpainting method^15^. Contrary to other models, Paired Cell Inpainting requires two channels for training - cytoplasmic background and target protein; hence we performed training of Paired Cell Inpainting using the GFP and cytoplasmic channels of the test images across 4,049 GFP-tagged proteins. We used the exact same architecture and training objective described by Lu et. al. The objective function minimizes a standard pixel-wise mean-squared error loss between the predicted target protein 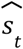and the actual target protein *s*_*t*_ (h and w are pixels across image width and height respectively):

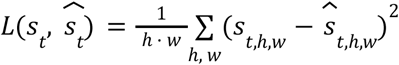

We performed hyperparameter optimilzation and selected an optimal learning rate of 1e-4 from {1e-4, 3e-4, 5e-4, 8e-4, 1e-3, 3e-3, 5e-3}. The Model was trained with Adam optimizer for 30 epochs (3 runs in total), and model weights were saved every epoch for subsequent evaluation. We selected the final model with early stopping based on the minimal validation set loss. After training, we extracted feature profiles of the test set images by maximum pooling the output of an intermediate convolutional layer, across spatial dimensions, as suggested by Lu et. al.^15^

### Evaluation of aFPs

Functional benchmarks used for assessment of the quality of the resulting feature profiles were derived from Gene Ontology Cellular Component^29^ (4045 protein annotations), Gene Ontology Slim Biological Process^29^ (3968 protein annotations), KEGG pathways^30^ (1422 protein annotations) and EMBL protein complexes^31^ (1402 protein annotations). Dubious ORFs and proteins without annotations were left out during comparison. To evaluate resulting features without further fine-tuning, we used strategies from two distinct perspectives: information retrieval and clustering quality.

Following standard practice, we computed pairwise distances across all available aFPs (4049×4049 distances in total) and sorted them from highest to lowest. Then protein pairs (which were not left out) were marked as positive if they had the same annotations, or negative otherwise, and AP and F-scores were computed. For proteins to be considered a positive pair, we required an exact agreement between labels in case of pathways and protein complexes standards, while for GO annotations we required at least 50% of the labels to overlap (due to high quantity of assigned labels). Results reported in **Fig. 2b, c** are based on ranking aFP pairs with correlation distance, and we found similar trends when using euclidean and cosine distances. We chose to continue analyses with the correlation metric due its lower susceptibility to fluctuations in individual feature values, and hence higher tolerance to outliers, which is a desirable property for the PIFiA workflow. For the clustering-driven benchmark, we clustered aFPs and compared clusters to the sets of proteins annotated to a certain term, and for each standard (we required cluster size to be at least 2 to be informative). For comparison, we applied AMI^34^ score between the resulting clusters and protein groups related to a certain term (with a higher score indicating more agreement between clusters and standard-defined categories). To obtain an AMI score for each method, we performed hierarchical clustering (with average linkage and correlation as a distance) of its per-protein representations and derived clusters across all similarity thresholds between 0.1 and 0.95 with a step of 0.05, and reported the maximal AMI across clusterings. For each deep learning model, feature profile evaluation was performed across 3 runs (results shown with bar plots in **Fig. 2**).

### Visualization of PIFiA feature profiles (aFPs and scFPs)

We used t-SNE^35^ for visualization of PIFiA feature profiles. We set the perplexity parameter to 40 for visualization of whole-proteome feature profiles averaged on the per-protein level (∼4000 points) (**Fig. 3c, d, f**), and to 200 for visualizing single-cell feature profiles from the test set (>100,000 points) (**Fig. 5a**). We represented the distribution of fundamental GO bioprocesses with a kernel-density estimate (KDE) using Gaussian kernels^36^ (**Fig. 3d**). We applied outlier filtering by removing points that do not lie within two standard deviations from the mean (across x or y t-SNE coordinates). We used Scott’s rule for KDE bandwidth selection^68^.

### Hierarchical clustering with aFPs

We performed agglomerative hierarchical clustering^33^ of the whole-proteome aFPs (4049 in total) using correlation as a distance metric and average linkage. The optimal cut-off distance for the whole-proteome hierarchical clustering was determined using the AMI curve between clustering labels and provided standard annotations, following the diminishing returns principle to find the elbow point. At the optimal distance cutoff, the slope of the curve becomes negligible, indicating that the available clusters cover most of the standard’s functional groups. In **Fig. 3** we used GO Cellular Component^29^ annotations as a standard and calculated clustering labels at different correlation thresholds between 0 and 1, with a step of 0.01. Clustering was performed on the whole-proteome feature profiles, and AMI scores were calculated on a subset of proteins that had a single annotation according to GO Cellular Component. We identified an optimal cut-off point when the derivative of the AMI curve (calculated over 20 steps, starting at correlation of 1) was less than a threshold of 0.1. The proposed strategy can be used on different standards, without requiring annotations to cover all proteins (**Fig. S2a**).

### Training logistic regression

To perform localization mapping, we trained a multinomial logistic regression (LR) using single-cell feature profiles obtained with PIFiA from the training set. We used supervised labels from 17 manually annotated localizations defined by Huh et. al.^10^ (we left out “ambiguous” category and classes with 5 or less proteins), and limited our training set to proteins that had a single annotated localization. Overall, our training set consisted of 1,432,774 single-cell feature profiles and included 2415 proteins from mitochondrion (465), nucleus (472), cytoplasm (799), actin (27), ER (245), vacuole (95), bud neck (8), spindle pole (35), Golgi (15), peroxisome (20), vacuolar membrane (47), cell periphery (51), nuclear periphery (45), endosome (28), and nucleolus (63). We followed our previously described dataset split (each protein’s single-cell crops were split into train, validation and test sets with 8:1:1 ratios). We used NVIDIA T4 GPU with 12GB of VRAM, and up to 16GB of system RAM on a single CPU to accelerate training; we trained LR for 5 epochs using Adam optimizer^63^ (1e-3 learning rate) and cross-entropy as a training loss; LR weights were saved after every epoch and we selected the final LR model with early stopping based on maximal validation set accuracy. Of note, after 2 epochs LR predictions stabilized and the difference between subsequent models was minimal (less than 3% test accuracy deviations).

To evaluate the quality of LR predictions, we compared its test set performance with DeepLoc^8^ (training procedure described in the benchmarking section). DeepLoc and LR were trained and evaluated on the sets of the same size and protein composition, with the only difference being DeepLoc used image crops for training, while LR using self-supervised scFPs from the pre-trained PIFiA model. Precision-recall curves for PIFiA LR and DeepLoc were generated on unseen scFPs and corresponding images from the test set of the single-localizing proteins (**Fig. S3b**).

### Sub-compartmental clustering with aFPs

We performed sub-compartmental clustering using single-localizing aFPs from the test set that were classified to the same localization by the previously described LR. For an aFP to be single-localizing, we required that its highest softmax probability was at least 0.6, and second-highest was no greater than 0.2 (more detailed analysis of single-localizing proteins and localization heterogeneity is described in the next section). We clustered aFPs of proteins that mapped to the same localization category and produced 15 per-compartment hierarchical trees (we used average linkage and correlation distance for clustering).

We calculated the Silhouette score^69^ using scikit-learn library, as the mean intra-cluster distance (a) and the mean nearest-cluster distance (b) for each aFP. The Silhouette coefficient for a sample is (b - a) / max(a, b); b is the distance between a sample and the nearest cluster that the sample is not a part of. We surveyed 15 per-localization hierarchical trees, clustered with average linkage and correlation metric, using correlation thresholds between 0.25 and 0.75, with a step of 0.05. Median of Silhouette scores across all localizations for a given threshold is shown in **Fig. S2e**. We found that thresholds between 0.5 and 0.6 yield maximal Silhouette scores, and a distance threshold of 0.5 corresponded to maximal GO bioprocess AMI value for whole-proteome aFPs clustering. Hence, we chose a 0.5 threshold to cluster aFPs belonging to each per-localization tree, and obtained 30 clusters, which we subsequently called sub-compartmental groups.

### Analysis of localization heterogeneity with scFPs

We used scFPs from the test set to analyse whole-proteome localization heterogeneity patterns. First, we used the pre-trained LR on 17 localization categories (described in the previous section) to map each protein’s test set scFPs to one of the 17 localization classes. We observed that the most probable localization class had an average 0.74 probability per protein (computed across all test set scFPs), while 2nd and 3rd classes scored 0.11 and 0.052 per-protein probabilities respectively. This motivated us to perform heterogeneity analysis with the two most probable localization categories, to avoid low scFPs quantities and potential noise effects. We then computed a mean probability across each localization class to determine the two most frequent localizations of the protein. Hence, for each protein *X* we obtained a distribution of 2-dimensional real valued probability vectors [*p*_*i*1_, *p*_*i*2_] with *p*_*i*1_ and *p*_*i*2_ corresponding to the probabilities of the first and second most frequent localization classes, *i*∈{1,…, *N*}, *N* is the number of test set scFPs of the protein *X*. Given this distribution, we could compute whether protein *X* is single-localizing or has AND-type / OR-type localization heterogeneity. We filtered low-confidence scFPs *i* whose sum of probabilities was below a confidence threshold: *p*_*i*1_ + *p*_*i*2_ < α_*conf*_ (low confidence region). Next, based on a heterogeneity threshold β, we divided the rest of the scFPs into first localization if *p*_*i*1_ > *p*_*i*2_ + β, second localization if *p*_*i*2_ > *p*_*i*1_ + β, or mixed-localizing category otherwise. We varied values of α_*conf*_ between 0.5 and 0.9, and values of β between 0.25 and 0.75 (with a step of 0.05), inspecting numbers of assignments into localization categories and low-confidence region, and selected α_*conf*_ and β as 0.5 based on elbow point analysis. Hence, scFPs of each protein were mapped into one of four classes - primary localization, secondary localization, mixed localization or low-confidence region (**Fig. 4d**). If we assume that percentages of the corresponding categories for protein *X* are *c*_1_, *c*_2_, *m, k*, then protein *X* would be marked as AND-type localizing if mixed category had higher percentage of scFPs than primary and secondary localizations together *m* > *c*_1_ + *c*_2_; otherwise protein *X* would be marked as OR-type if no less than 8% of scFPs belonged to the secondary localization *min*(1, *c*_2_) > 0. 08, and single-localizing in the other case. We experimented with OR-type thresholds between 0.05 and 0.3 (with a step of 0.01), and found that the number of OR-type localizing proteins saturated between 0.07 and 0.1 thresholds. We selected 0.08 as an elbow point between *min*(1, *c*_2_) value and number of category assignments. Thus, each protein was marked as single-localizing, OR-type, AND-type or undetermined (if too many scFPs were assigned as low-confidence).

### Cell cycle prediction and annotation with scFPs

We trained an ensemble of three CNNs for cell cycle classification using cytosolic and nuclear channels from our dataset. These two channels contained enough information to distinguish the cell cycle stage of a cell. The CNN contains four convolutional blocks followed by two fully-connected layers, and was trained to predict one of four cell cycle stages - G1, S, metaphase and anaphase (MA), or telophase (T) (**Fig. S6a**).

We manually labeled 800 crops of cells from 103 different proteins, corresponding to distinct cell cycle stages (with 200 crops from each class) and used heavy data augmentation during training to prevent overfitting:rotation by arbitrary angle, vertical and horizontal flips, image zoom within 0.02 range, and vertical and horizontal shifts of up to 9 pixels. We used three-fold cross-validation. The training was performed on 64×64×2 dimensional crops over 150 epochs using Adam optimizer, with loss being a categorical cross-entropy across 4 cell cycle categories. We performed hyperparameter optimization to select learning rate (5e-4) and dropout rate (0.05). We performed 3 independent runs with random weights initialization (using truncated normal distribution with a standard deviation of 0.1). Model weights were saved after every epoch, and final models for each run were selected with early stopping based on maximal validation accuracy. Training and test accuracy and categorical cross-entropy loss are shown in **Fig. S6b**. We created an ensemble of three CNNs (from epochs corresponding to minimal loss value), and subsequently mapped each single-cell crop to a 4-dimensional real valued vector of cell cycle probabilities. The cell cycle probability vector was computed as an average of three CNNs’ probability vectors of that crop. We subsequently joined T and G1 categories due to high cell density in certain crops, which could potentially lead to an incorrect cell cycle category assignment.

We applied Mann-Whitney U test^48^ to identify proteins whose localization changes had cell cycle dependency. For each protein with localization heterogeneity, we annotated its single-cell crops from train, validation and test sets using both LR localization categories and cell cycle stages (via cell cycle classifier). We selected two primary annotated localizations, and compared cell cycle stage distribution of the corresponding crops. For each cell cycle stage, our null hypothesis was that the stage was equally represented among both localizations. Localizations with significant distribution differences (i.e. p-value < 1e-3) were annotated as related to the particular cell cycle stage.

### Functional enrichment analysis

Gene Ontology (GO) enrichments were performed using GO-term Finder Version 0.86, available through the *Saccharomyces cerevisiae* Genome Database (https://www.yeastgenome.org/goTermFinder). We applied gene set enrichment analysis (GSEA) using Python package GSEApy^70^ (https://github.com/zqfang/GSEApy) to analyse hierarchically clustered protein groups, sub-compartmental groups and sets of multi-localizing and mixed-localizing proteins (**Table S2, S3**). Query gene sets for GSEA included GO biological process, cellular component and molecular function standards^29^. GSEA results were filtered to include gene sets with p-values below 0.05 and a minimum gene set size of 2. We applied Bonferroni correction to obtain adjusted p-values. We also applied one-sided Fisher’s exact test with Costanzo group 19^32^ categories to analyse nucleus OR cytoplasm, nucleus AND cytoplasm gene sets, reporting protein sets with p-value < 0.05 as the ones showing enrichment (**Table S3**).

### Discovery of interacting proteins from scFPs

We identified clusters containing potentially interacting proteins using two steps. First, we hierarchically clustered scFPs from the test set (we used average linkage and a correlation metric). After that, we divided the dendrogram from top to bottom and traced the number of unique proteins inside the cluster along with the division thresholds of 0.05 points. We found thresholds of the dendrogram at which the number of proteins in a cluster plateaus (95% of protein composition remains the same). After such “morphologically inseparable” clusters were identified, we used three data-driven scores to measure the quality of the resulting clusters - cell ratio, elbow point, and child ratio. Cell ratio *c* is an average percentage of a protein’s cells that fall into a particular cluster. A higher cell ratio translates into less dispersed cells of the same protein, and more confident protein assignment into the particular cluster. Elbow point *k* is a clustering distance at the level of the current cut (1-PCC in our case). A lower elbow point corresponds to a smaller distance between proteins in their feature profiles space. Descendant ratio *d* of the particular root cluster is the percentage of its descendent clusters that were annotated to the same root. A high child ratio corresponds to more agreement of the child clusters, hence indicating a more confident prediction. We devise a final score S as follows:

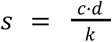

We used a final score cutoff of 0.6 to produce a list of 88 high-confidence clusters. We compared performance of our adaptive thresholding approach with clustering approaches from three different families - connectivity (hierarchical clustering^33^), centroid (k-means^49^) and density methods (DBSCAN^50^) (**Fig. 5c**). We performed clustering on the same test set of scFPs with different approaches. For each of the methods used in our comparison, we tried a range of hyperparameters (k ranging from 5 to 500 with a step of 5 in k-means, epsilon ranging from 0.1 to 5 with a step of 0.1 in DBSCAN, and correlation threshold ranging from 0.05 to 0.5 with a step of 0.025 for hierarchical clustering) and report the ones corresponding to the maximal median F1-score across all clusters. F1 scores were calculated by assigning each pair of scFPs ground truth label (0 or 1 depending on whether they are part of the same protein complex) and predicted label (0 or 1 depending on whether they are part of the same cluster).

### Gradient maps

We applied the SmoothGrad method to obtain per-feature gradient maps of the input images^55^. Original gradient maps *m*_*c*_ (*x*) compute the derivative of activation function *S* of the highest-scoring class *c* with respect to the input image *x*, and thus highlight pixels which influence classification decision:

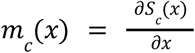

Since we were interested in feature interpretation, but not interpreting the classification decision, we modified this computation. In our implementation of gradient map *M*_*i*_ (*x*), we take the derivative of specific feature *f*_*i*_ from the feature vector *f* with respect to the input image *x*:

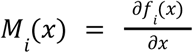

Hence, our gradient maps highlight regions of the image that impact the value of the selected feature. SmoothGrad produces a gradient map *M*_*i*_ (*x*) by averaging a number of gradient maps obtained from an input image with added noise *N*(0, σ^2^):

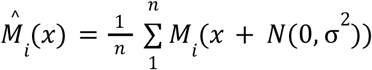

We used n=100 images and σ=0.05 noise level.

## Data availability

The image data used in this work are available at https://thecellvision.org/pifia/.

## Code availability

Source code for the PIFiA network and downstream analysis is available at https://github.com/arazd/pifia.

## Acknowledgements

We thank Oren Kraus, Michael Costanzo, Nil Sahin, Alan Moses, Leah Cowen and Matej Usaj for valuable discussions and advice. This work was supported by grants from the National Institutes of Health (R01HG005853 to B.A., C.B.), and the Canadian Institutes of Health Research (PJT-180259 to B.A.). Equipment for automated image acquisition and analysis was purchased using funds from the Canadian Foundation for Innovation and the Ontario Research Fund. J.B. was supported by the Canadian Institute for Advanced Research (CIFAR) AI Chairs program and the National Sciences and Engineering Research Council (Canada). Computational resources were provided, in part, by the Province of Ontario and the Government of Canada through the Vector Institute for Artificial Intelligence. A.R. was supported by the Province of Ontario (Ontario Graduate Scholarship, 2021-2022) and the Vector Institute for Artificial Intelligence (Vector Institute Postgraduate Affiliate Scholarship, 2019-2021). C.B. is a Fellow of the CIFAR.

## Author contributions

A.R. developed the PIFiA pipeline and performed the computational experiments. A.B. designed components of the downstream data analysis. M.M.U. and H.F. performed biological analysis and interpretation of the derived clusters. M.P.D.M. developed the PIFiA visualization tool for the CellVision website. H.G.S. constructed and imaged the modified yeast GFP collection. K.W. performed the co-localization experiments. A.R., H.F., A.B., M.M.U., C.B., J.B. and B.A. wrote the manuscript. B.A., C.B. and J.B. conceived and supervised the project.

## Supplementary Figures

**Sup. Figure 1.**
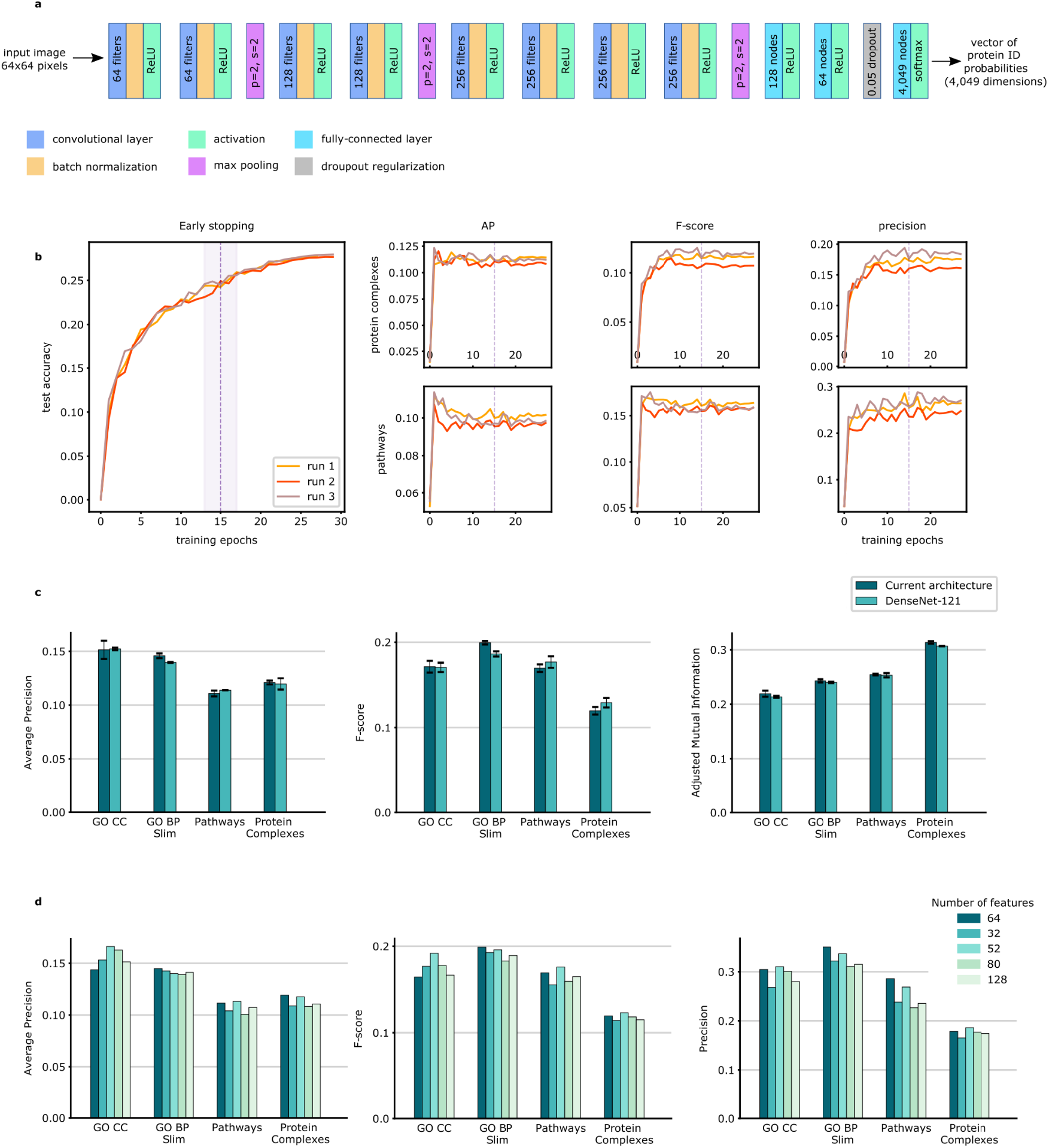
PIFiA network architecture and training settings. **a**, Overview of the architecture of PIFiA convolutional network. **b**, Left plot: test accuracies of three different runs over the course of training (X axis: epochs, Y axis: test accuracy). Smaller plots: average precision, F-score and precision on protein complexes and pathways standards (X axis: epochs, Y axis: corresponding score on test set). The purple line indicates point of early stopping, when accuracy starts to saturate (derivative of the test accuracy smaller than a threshold of 0.5%). **c**, Comparison of the current PIFiA architecture with a common baseline, DenseNet-121, across four different standards (Gene Ontology Cellular Component, Gene Ontology Bioprocess Slim, KEGG Pathways, EBI Protein complexes) in terms of average precision, F-score and adjusted mutual information. **d**, PIFiA performance across different dimensions of the feature profiles (32, 52, 64, 80, 128).

**Sup. Figure 2.**
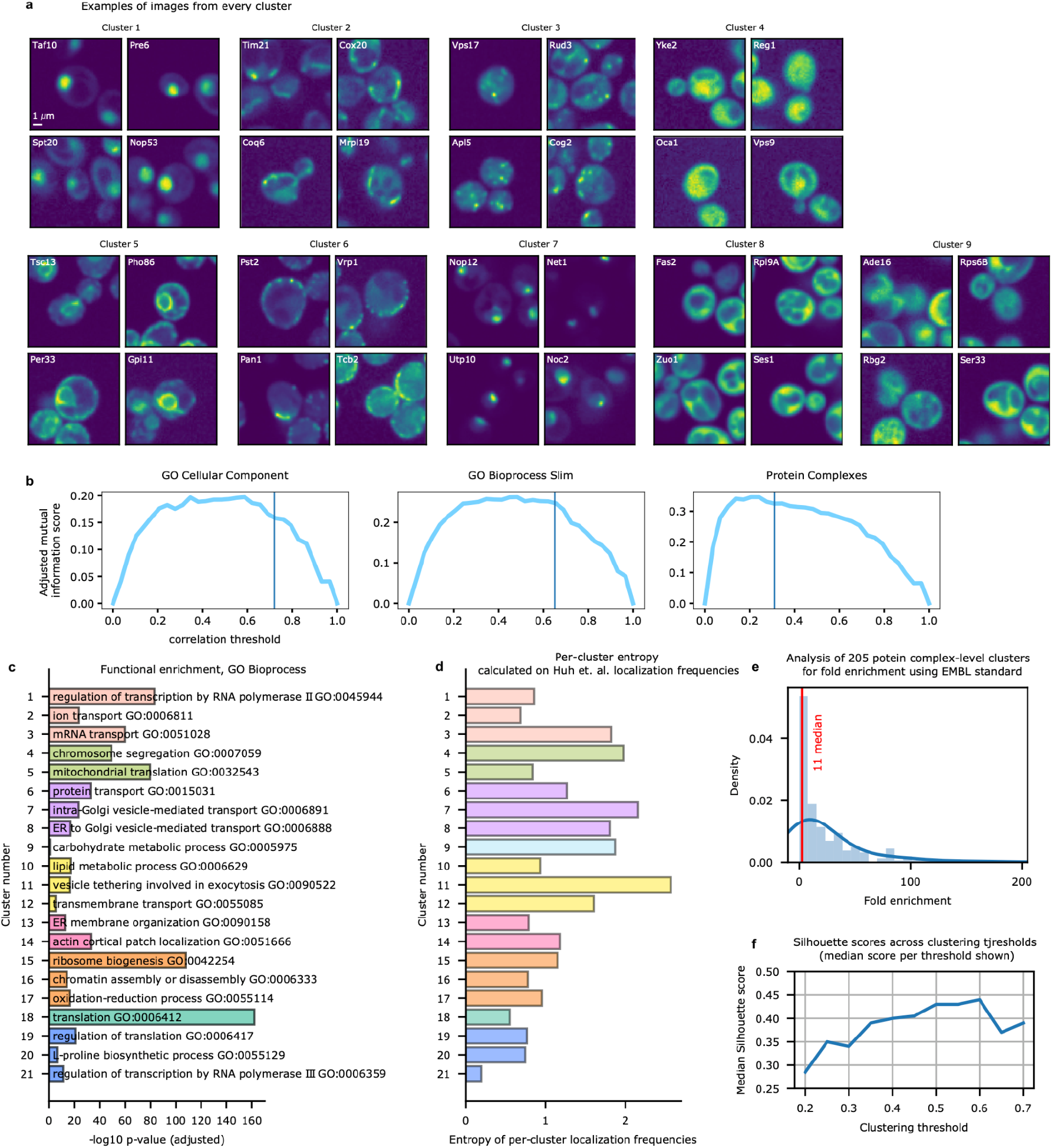
Examples of AMI cutoffs and cell images from nine clusters. **a**, Examples of GFP-tagged proteins from nine clusters corresponding to major cellular components. **b**, Adjusted mutual information across different correlation thresholds for cellular component, bioprocess and protein complex standards (X axis: correlation threshold, Y axis: adjusted mutual information score). Vertical line indicates a point of a dendrogram cut determined by saturation of a score. **c**,The top Gene Ontology Biological Process terms and corresponding functional enrichments are shown for 21 clusters obtained from clustering by GO Biological Process Slim cutoff; color coding corresponds to parent cell component clusters defined in Fig. 3a, b. **d**, Entropy across 21 bioprocess clusters calculated from localization category frequencies (from Huh et al. (2003) standard). **e**, Distribution of fold enrichments on EMBL protein complex standard for 205 clusters derived from Protein Complex cutoff. f, Silhouette scores for different thresholds across single-localizing aFPs from the same localization category (X axis: clustering thresholds, Y axis: median Silhouette score across all localizations).

**Sup. Figure 3.**
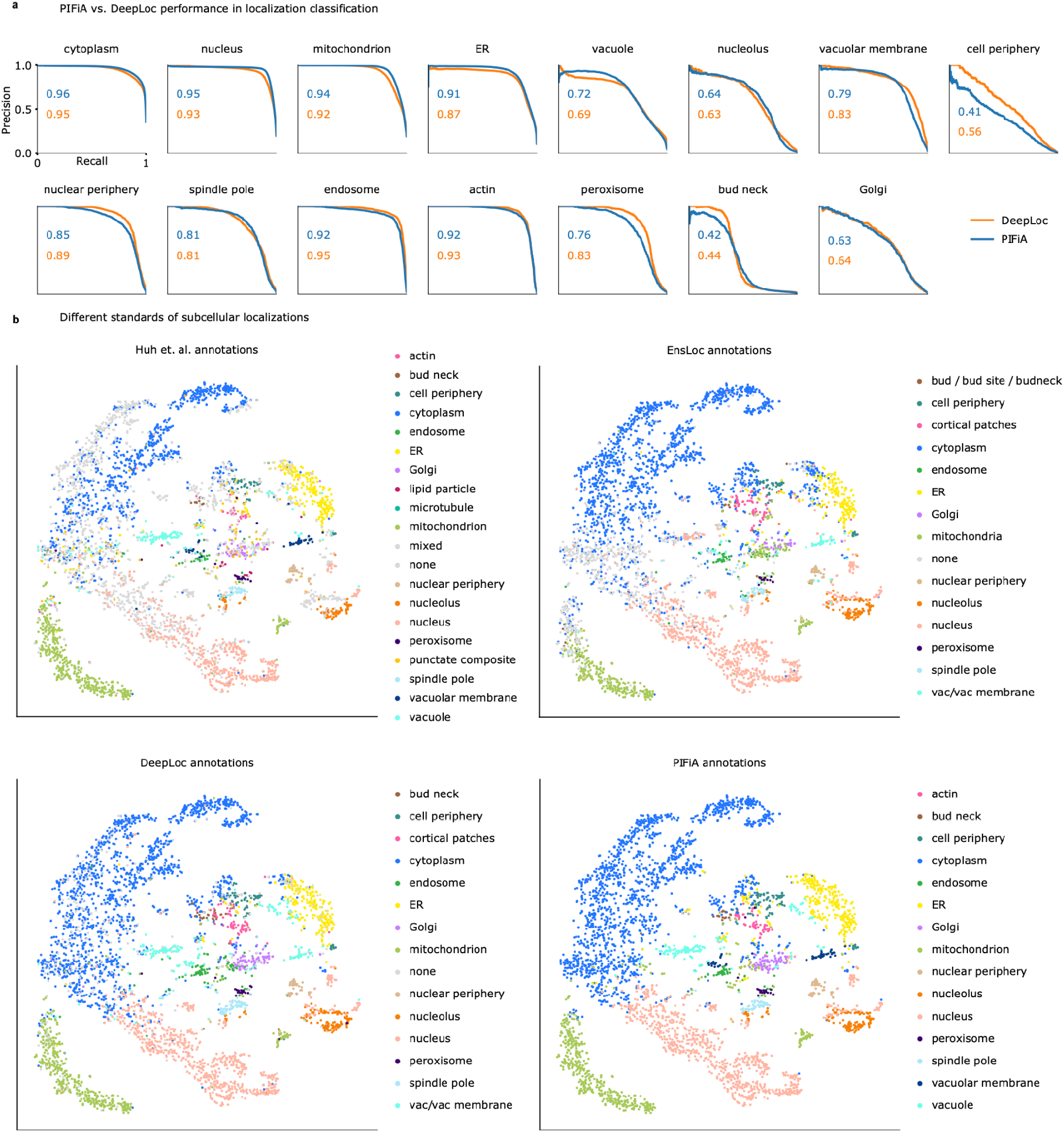
Comparison of PIFiA annotations and existing localization standards. **a**, Comparison of localization classification performance of DeepLoc versus PIFiA feature profiles coupled with a logistic regression. Precision-recall plots are shown for 15 subcellular localizations (X axis: recall, Y axis: precision). **b**, Whole-proteome aFPs tSNE colored by different annotations of subcellular localization: manual annotations from Huh et al. (2003), and computationally-derived annotations from EnsLoc, DeepLoc and PIFiA.

**Sup. Figure 4.**
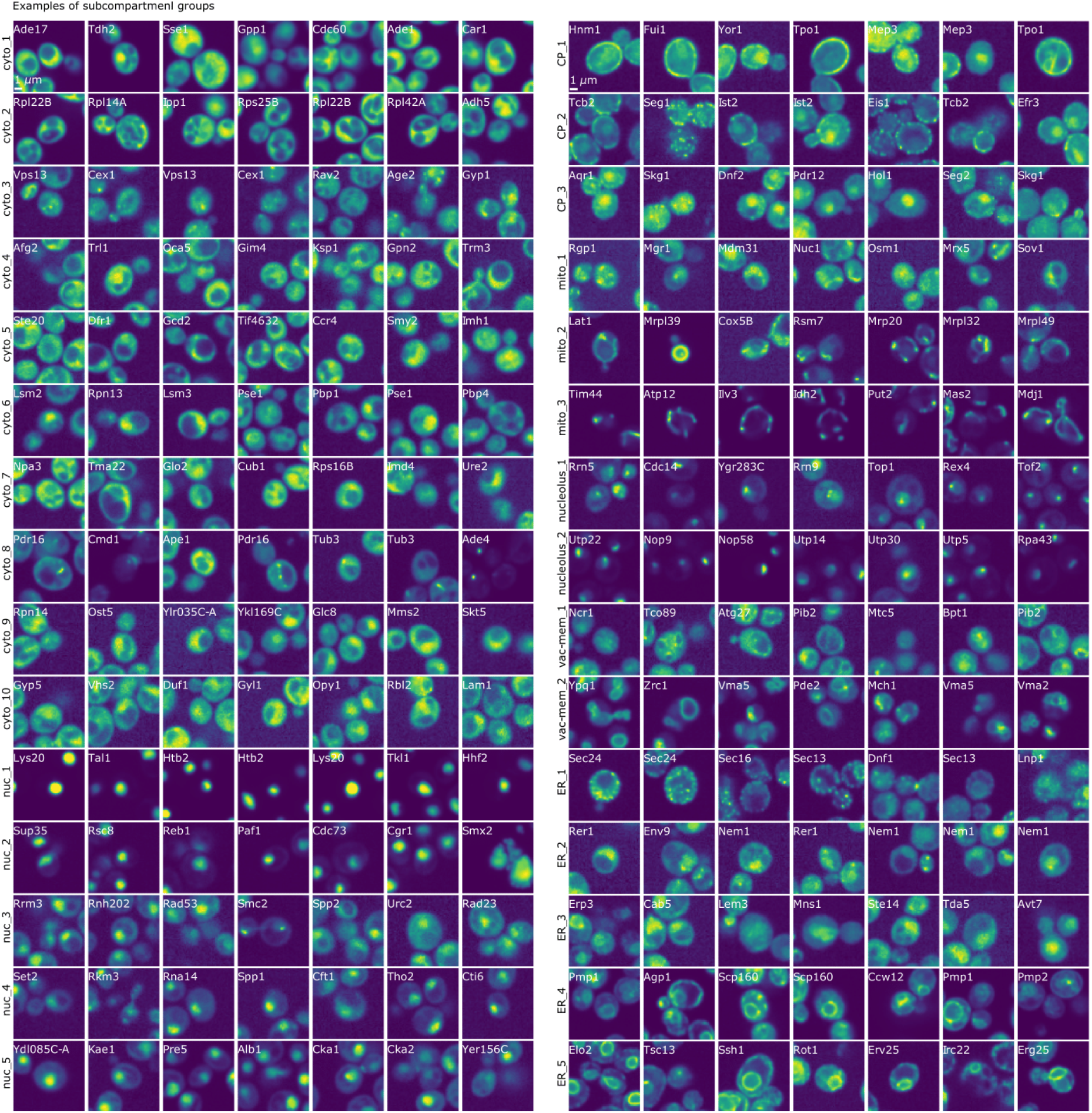
Examples of proteins from different sub-compartmental groups. Examples of GFP-tagged proteins from 30 sub-compartmental groups. Each row corresponds to a sub-compartmental cluster (e.g. nuc-1, nuc-2). The relevant GFP-tagged protein is identified on each micrograph.

**Sup. Figure 5.**
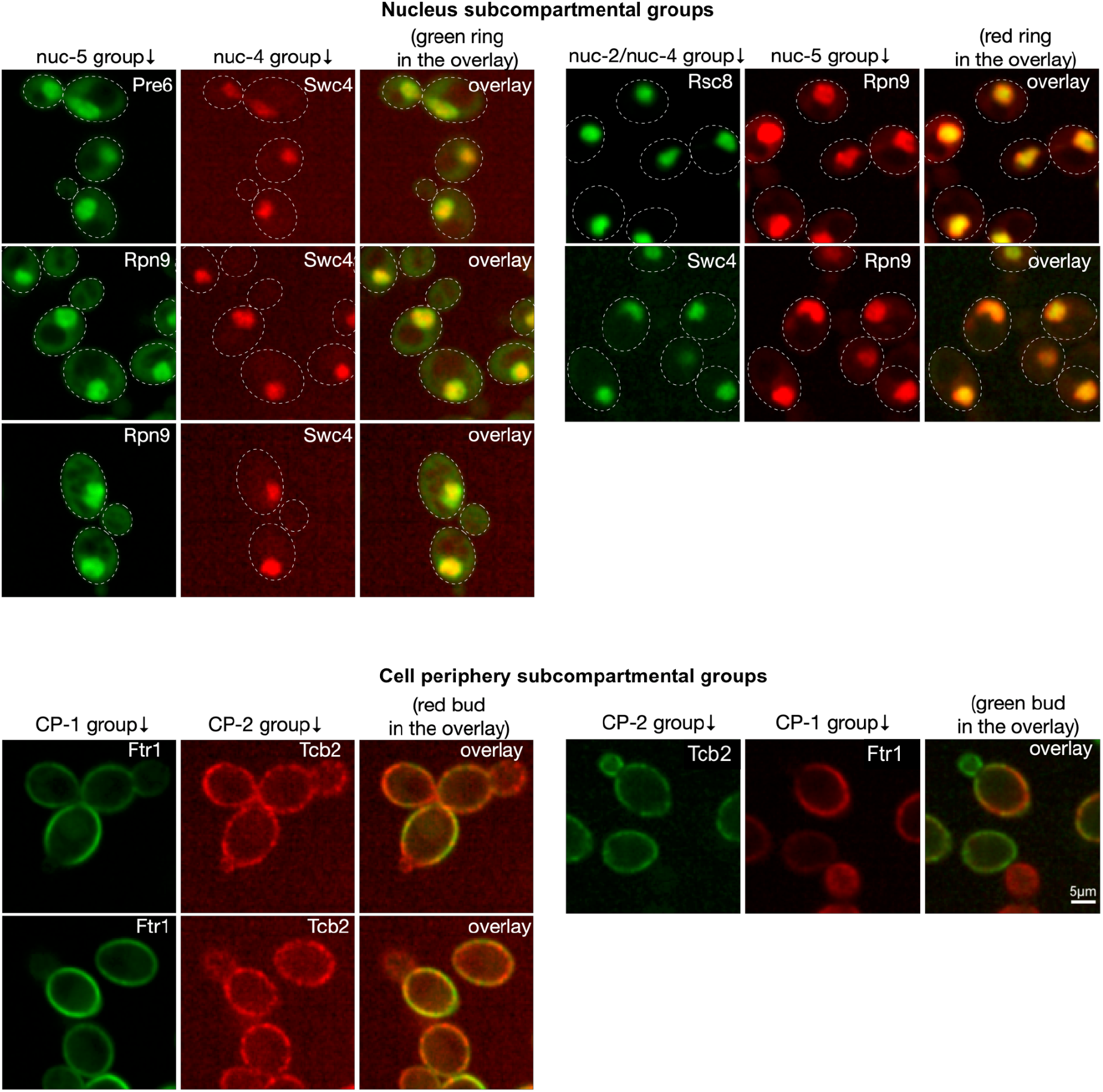
Examples of proteins from different sub-compartmental groups. Colocalization experiment results: representative micrographs of cells expressing mNeonGreen- (green images) or mScarlet- (red images) tagged proteins annotated to nucleus (top panel) or cell periphery (bottom panel) groups. Overlays of the mNeonGreen and mScarlet images are shown on the right of each triplet of images. The tagged proteins are indicated on the micrographs (scale bar shown bottom right).

**Sup. Figure 6.**
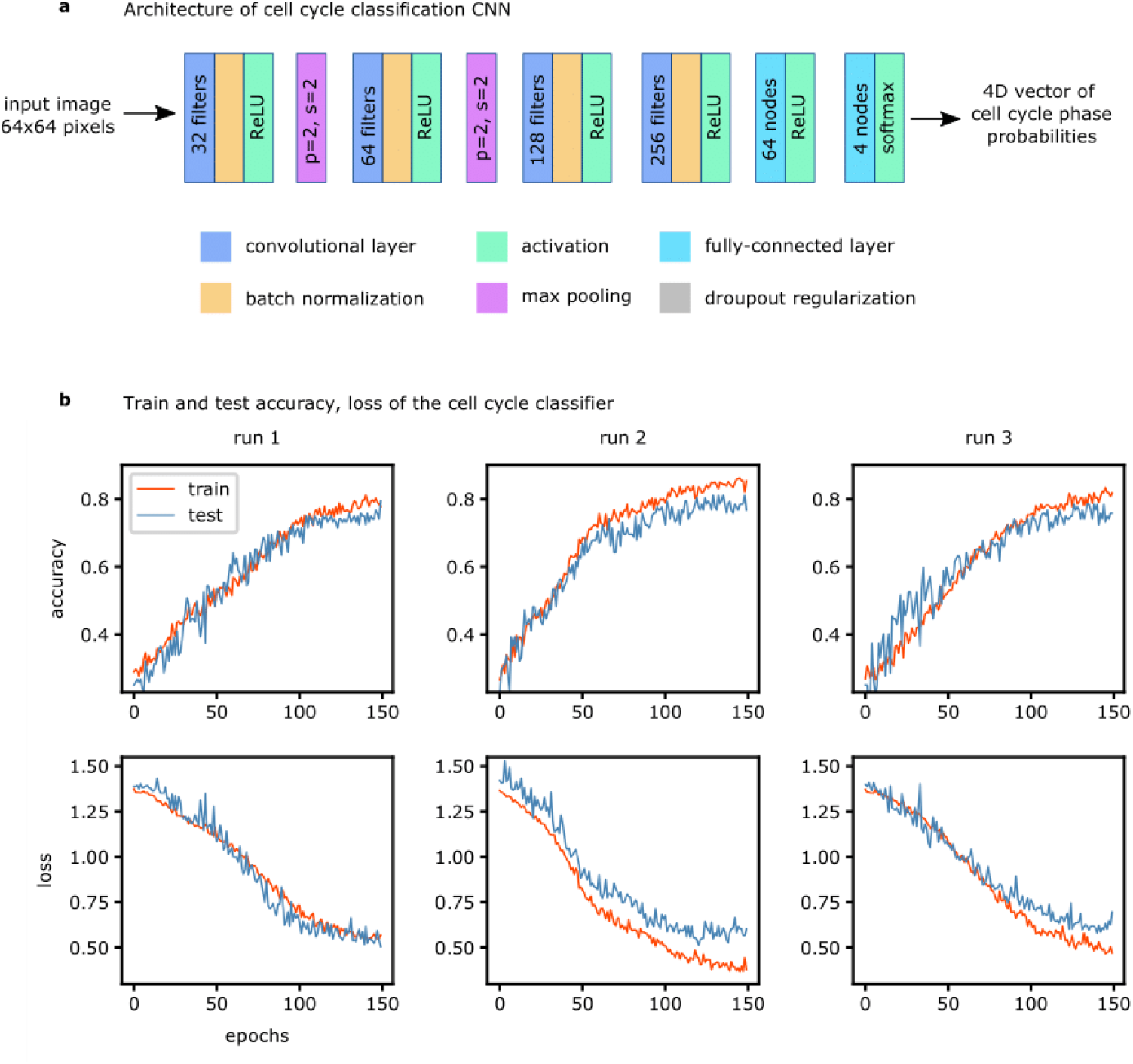
Cell cycle classifier training settings. **a**, Architecture of the convolutional neural network used for cell cycle classification. **b**, Train and test set performance (accuracy and loss) of the cell cycle classifier across three independent network runs.

**Sup. Figure 7.**
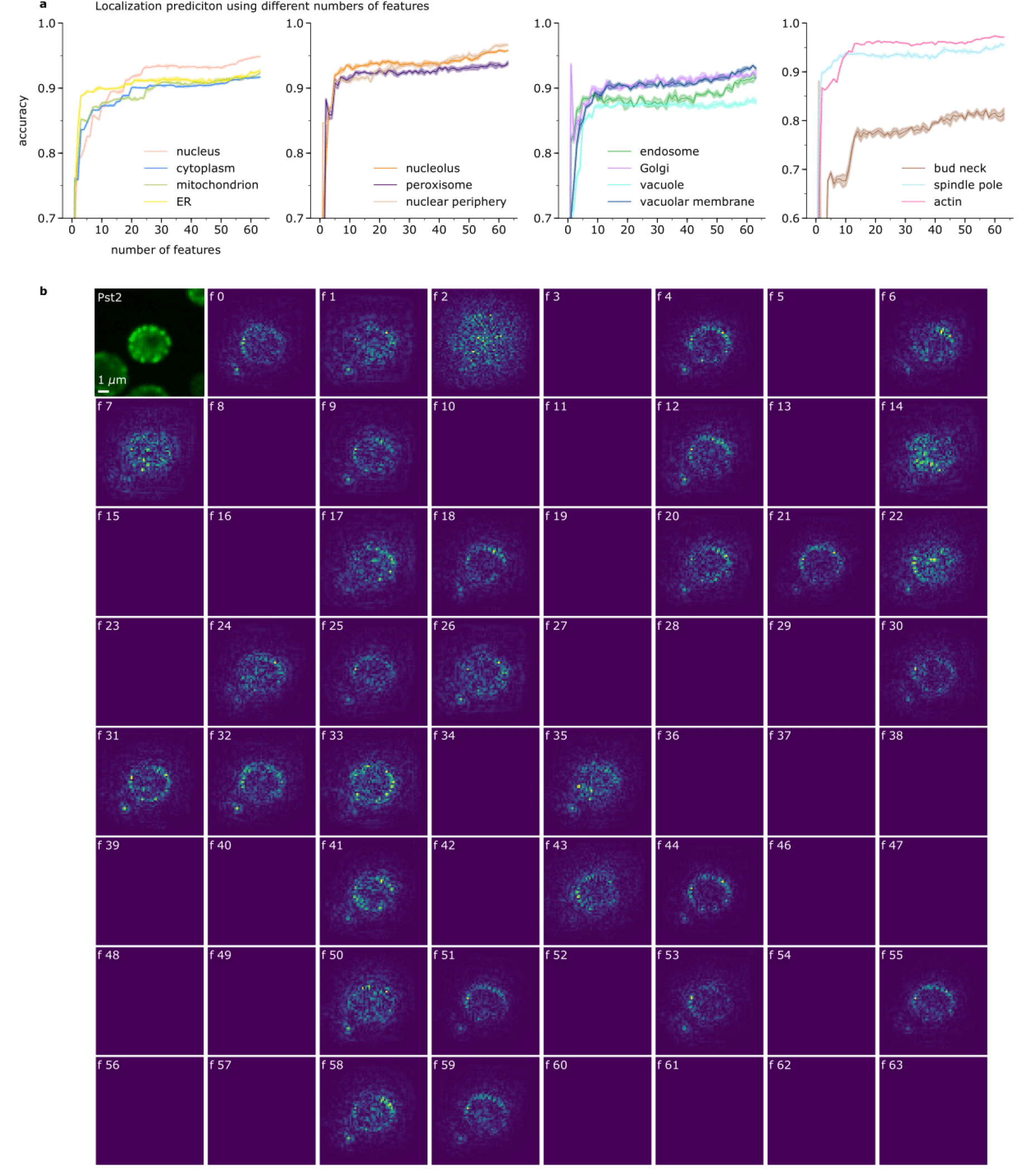
Interpretability studies. **a**, Accuracy of subcellular localization prediction using different dimensionality of feature profiles (X axis: number of features in a feature profile, Y axis: accuracy). **b**, Examples of feature-wise gradient maps obtained with SmoothGrad for all features of Pst2-GFP protein crop.

## Supplementary Tables

Supplementary tables are available at the link https://drive.google.com/drive/folders/19DAPPJeYeqovk6jRo_su0OX7tWYoAEUh?usp=sharing.

